# Protein interaction mapping reveals widespread targeting of development-related host transcription factors by phytoplasma effectors

**DOI:** 10.1101/2020.02.13.946517

**Authors:** Miguel Correa Marrero, Sylvain Capdevielle, Weijie Huang, Ali M. Al-Subhi, Marco Busscher, Jacqueline Busscher-Lange, Froukje van der Wal, Dick de Ridder, Aalt D.J. van Dijk, Saskia A. Hogenhout, Richard G.H. Immink

## Abstract

Phytoplasmas are pathogenic bacteria that reprogram plant host development for their own benefit. Previous studies have characterized a few different phytoplasma effector proteins that destabilize specific plant transcription factors. However, these are only a small fraction of the potential effectors used by phytoplasmas; therefore, the molecular mechanisms through which phytoplasmas modulate their hosts require further investigation. To obtain further insights into the phytoplasma infection mechanisms, we generated a protein-protein interaction network between a broad set of phytoplasma effectors and a large, unbiased collection of *Arabidopsis thaliana* transcription factors and transcriptional regulators. We found widespread, but specific, interactions between phytoplasma effectors and host transcription factors, especially those related to host developmental processes. In particular, many unrelated effectors target specific sets of TCP transcription factors, which regulate plant development and immunity. Comparison with other host-pathogen protein interaction networks shows that phytoplasma effectors have unusual targets, indicating that phytoplasmas have evolved a unique and unusual infection strategy. This study contributes a rich and solid data source that guides further investigations of the functions of individual effectors, as demonstrated for some herein. Moreover, the dataset provides insights into the underlying molecular mechanisms of phytoplasma infection.

## 1 Significance statement

This work shows that the effectors of phytoplasma, a globally economically important bacterial plant pathogen, have pervasive interactions with development-related host transcription factors, providing a way to take over plant growth and development in favor of the pathogen and its insect vector. The obtained comprehensive protein interaction network and a showcase of the potential biological consequences of a selected effector advance our understanding of phytoplasma-host interactions and provide guidance for further study.

## 2 Introduction

Among the vast number of plant-pathogenic microorganisms, some have the intriguing ability to manipulate the development of their hosts in order to increase their own fitness. A remarkable, well-known example is that of the rust fungus *Puccinia monoica*, which alters the morphology of its host to create pseudoflowers covered by fungal spermatogonia. These pseudoflowers successfully attract pollinators, which then spread the fungal reproductive cells [Roy, 1993]. The complex and diverse molecular underpinnings through which pathogens modify host development have been studied in recent years [Le Fevre et al., 2015], but we are far from a complete understanding of most mechanisms.

Phytoplasmas (*Candidatus* Phytoplasma), which are obligate bacterial plant pathogens, represent another example of a pathogen partially taking over plant development, and manipulating development of their plant hosts via processes that are yet to be fully understood. Phytoplasmas have a life cycle that alternates between plants and specific insect herbivores, such as leafhoppers, which transmit these bacteria to new plant hosts [Weintraub and Beanland, 2006]. The insects become phytoplasma carriers by acquiring these bacteria from the phloem of infected plants, and once the bacteria colonize the salivary glands of their insect vectors, they can be introduced into the phloem of new plant hosts when the insects feed [Sugio et al., 2011b, MacLean et al., 2011]. Phytoplasma-infected plants often display altered morphology [Bertaccini, 2007] that suggest interference with fundamental developmental processes. For example, the Aster Yellows phytoplasma strain Witches’ Broom (*Ca.* Phytoplasma asteris; AY-WB phytoplasma) induces phyllody (conversion of flowers into leaflike structures), virescence (green coloration of non-green floral tissue) and witches’ brooms (increased proliferation of stems, branches and leaves), and promotes attraction of AY-WB leafhopper vectors to infected plants [Sugio et al., 2011b,a, MacLean et al., 2014, Orlovskis and Hogenhout, 2016, Pecher et al., 2019, Huang et al., 2021, 2020, Huang and Hogenhout, 2022]. Additionally, infected plants are often sterile, and thus serve the sole purpose of feeding and propagating the bacteria.

While this phenomenon alone makes phytoplasmas very interesting pathogens, they are not merely a scientific curiosity. Phytoplasma infections are of socioeconomic importance; these obligate bacteria form a large and diverse group that began diversifying ±316 Mya, coinciding with the origins of seed plants and their sap-feeding insect vectors of the order Hemiptera [Cao et al., 2020] and are found in most vascular plant species, often but not always causing dramatic yield losses of crops and ornamental plants [Strauss, 2009]. Plants of high economic importance that suffer regular yield losses because of persistent phytoplasma outbreaks include lime [Donkersley et al., 2018], grape [Malembic-Maher et al., 2020], apple [Fránová et al., 2019] and coconut [Gurr et al., 2016]. More erratic outbreaks occur in herbaceous crops, such as carrot [Frost et al., 2013], maize [Jović et al., 2009] and tomatoes [Santos-Cervantes et al., 2008], as well as in ornamental plants and trees, such as flower bulbs, ash and elm [Cortés-Martínez et al., 2008, Herath et al., 2010, Sinclair et al., 2000]. Several phytoplasmas are considered quarantine pests [, PLH].

Plant pathogens produce effectors, proteins to hijack the host cell for improving the pathogen’s ftness in the host by different means (e.g. by modulating the host immune response) [Mattoo et al., 2007]. Four different phytoplasma effectors, SAP54, SAP05, SAP11, and TENGU, have been characterized, shedding light on the molecular mechanisms through which phytoplasmas manipulate their hosts. SAP54 mediates degradation of MIKC MADS-box transcription factors involved in floral development, inducing leaf-like flowers that resemble phyllody symptoms of phytoplasma-infected plants [MacLean et al., 2014], whereas SAP05 mediates degradation of both SBPs and GATA transcription factors, thereby prolonging the host lifespan and inducing witches’ broom-like proliferations of leaf and sterile shoots [Huang et al., 2021]. SAP11 binds and destabilizes certain TCP transcription factors involved in axillary meristem outgrowth, leaf shape determination and jasmonate signalling, leading to an altered morphology and decreased plant defense [Sugio et al., 2011a, 2014, Chang et al., 2018, Wang et al., 2018, Pecher et al., 2019]. These three SAP effectors promote attraction and reproduction of leafhopper vectors on plants [Sugio et al., 2011a,b, Orlovskis and Hogenhout, 2016, Huang and Hogenhout, 2022].Finally, TENGU causes dwarfism, witches’ broom and arrested flower development. Although its molecular mode-of-action has not been clearly identified, this effector downregulates expression of auxin response factors ARF6 and ARF8, leading to decreased jasmonate biosynthesis [Hoshi et al., 2009, Minato et al., 2014]. These alterations caused by individual effectors reflect the symptoms caused by phytoplasma infections, and, overall, suggest a trend of targeting plant transcription factors in order to exploit the host.

However, these four effectors are only a small part of the potential effector arsenal of phytoplasmas. Sequencing of phytoplasma genomes has allowed the identification of many more candidate effectors that the pathogen might use to colonize its hosts [Bai et al., 2009], some of which are preferentially expressed in the plant and others in the insect [MacLean et al., 2011, Oshima et al., 2011]. Given the precedent of targeting host transcription factors in order to manipulate plant development, we asked how prevalent this phenomenon is across phytoplasma effectors expressed in plants. In order to identify interactions between effectors and host transcription factors, we performed large-scale yeast-two hybrid (Y2H) assays, testing interactions between 21 phytoplasma effectors and a comprehensive library of *Arabidopsis thaliana* transcription factors and transcriptional regulators [Pruneda-Paz et al., 2014]. Such an approach is useful to obtain a systems-level view of host-pathogen interactions [Tripathi et al., 2019, Rodriguez et al., 2019].

Overall, the resulting protein-protein interaction (PPI) network shows pervasive interactions of candidate phytoplasma effectors with plant host transcription factors, particularly with those that regulate plant development. Furthermore, we find that numerous unrelated effectors interact with multiple TCP transcription factors, known to be targeted by diverse effector proteins of bacterial, fungal and oomycete pathogens [Mukhtar et al., 2011, Weßling et al., 2014]. Compared to other pathogens’ effectors, those of phytoplasmas target plant growth and development more intensely.

## 3 Material and methods

### 3.1 Yeast-two hybrid assays

The sequences corresponding to the mature versions (without signal peptides as predicted by [Bai et al., 2009]) of 21 selected phytoplasma effector proteins were amplified from the genomic DNA of plants infected by aster yellows witches’ broom phytoplasma (AY-WB) and cloned into pDONR207. Obtained fragments were sequenced for verification. The SAP11 homolog of maize bushy stunt phytoplasma (*Ca.* Phytoplasma asteris; MBSP) was amplified from MBSP-infected maize plants [Orlovskis et al., 2017]. Phytoplasma SAP54 genes from rapeseed phyllody phytoplasma (RP) and peanut witches’ broom phytoplasma (PnWB) and SAP11 from various phytoplasmas were identified from phytoplasma genome sequences in the public domain, and the sequences corresponding to the mature protein without signal peptide were synthesized (General Biosystems, North Carolina, USA) and cloned into pDONR207. The effector DNAs were sub-cloned from pDONR207 into the Y2H bait vector pDEST32 by Gateway-based recombination. The resulting effector bait plasmids were transformed into yeast strain PJ69-4 mating type Alpha, followed by a test for autoactivation of reporter genes, as described previously [de Folter and Immink, 2011]. Subsequently, matrix-based Y2H screenings were performed following the protocol in [de Folter and Immink, 2011]. Each bait was screened against the *Arabidopsis* transcription factor collection, consisting of 1980 clones [Pruneda-Paz et al., 2014]. For the baits that did not exhibit autoactivation, incubation was done on SD medium lacking Leucine, Tryptophan, and Histidine and supplemented with either 1 or 5 mM 3-amino-1,2,4-triazole (3-AT), respectively. For SAP06 and SAP48, which showed autoactivation up to 15 mM 3-AT, the incubation was done on SD medium lacking Leucine, Tryptophan, and Histidine and supplemented with either 20 or 25 mM 3-amino-1,2,4-triazole (3-AT), or SD medium lacking Leucine, Tryptophan, and Adenine, respectively. Yeast growth and hence, protein interaction events were scored after incubation of yeast at 20 °C for 6 days. The interactions are provided in MIAPE (Minimum Information About a Proteomics Experiment) compliant format (Table S1).

For further analyses of SAP11 interactions with TCP transcription factors, sequences corresponding to the mature proteins (without signal peptides) of various SAP11 homologs or SAP11 chimeras were synthesized by General Biosystems (North Carolina, USA) and subcloned into the pDESTGBKT7 vector (Tables S2 & S3). Representative TCP genes were cloned into the pDESTGADT7 vector. These plasmids were transformed into the yeast strain AH109 for testing protein-protein interaction using the Matchmaker Gold system (Takara Bio, Kusatsu, Japan). Yeast transformation and growth were performed as described in [Pecher et al., 2019], and the SAP11-TCP interactions were studied in at least three independent experiments.

### 3.2 BiFC in Nicotiana benthamiana

BiFC constructs were generated by Gateway cloning of the full-length ORFs of the *Arabidopsis* genes encoding the putative SAP06 interactors into the pDEST-SCYCE(R)GW vector and the synthetic plant codon-optimized ORF of the mature SAP06 protein (without signal peptide) into the pDEST-SCYNE(R)GW vector [Gehl et al., 2009]. For this purpose, the coding sequences of SAP06, GAI (AT1G14920), RGA1 (AT2G01570), RGL1 (AT1G66350), and RGL3 (AT5G17490) in the pDONR207 vector were used in the Gateway LR reaction. Leaves from four-week-old *Nicotiana benthamiana* plants were transiently infected as described previously [Diaz-Granados et al., 2020], followed by confocal imaging to check for cyan fluorescent protein signal three days after infection. As positive control, the combination SCYCE-GpRBP1 + SCYNE-StUPL3 was included [Diaz-Granados et al., 2020], and as negative control the combination SCYCE-GpRBP1 + SCYNE-SAP06.

### 3.3 Host protein family annotation

We assigned each host protein to a particular family by first checking PlantTFDB, a comprehensive database of plant transcription factors [Jin et al., 2017]. However, not all host proteins could be found in this database. The host library is based on proteins from different resources [Pruneda-Paz et al., 2014], including PlnTFDB [Pérez-Rodŕıguez et al., 2010], which is more general and contains not only transcription factors, but also other transcriptional regulators. Therefore, we used PlnTFDB to assign families to each host protein absent in PlantTFDB. In total, we were able to assign 1830 of the 1980 proteins to a unique family (1434 found in PlantTFDB and 396 in PlnTFDB). The remaining proteins are marked as unannotated.

### 3.4 Statistical analysis of the phytoplasma-*Arabidopsis* PPI network

Gene Ontology (GO) enrichment analyses were performed using goatools v0.8.4 [Klopfenstein et al., 2018] with the host protein library as background. Genes annotated with a GO term were automatically annotated with the parent terms as well, and multiple testing correction was performed with the Benjamini-Hochberg (BH) method (false discovery rate threshold: 0.05) [Benjamini and Hochberg, 1995]. 98% of the proteins in the host library were present in the GO annotation file. All of the analyses excluded terms inferred from electronic annotation (i.e. those with the evidence code IEA).

As there might be an overall trend of effectors interacting predominantly with specific host protein families, we tested whether each host protein family is enriched in the PPI network with respect to the host protein library. We did this using Fisher’s exact test. We performed multiple testing correction using the BH method (false discovery rate threshold: 0.05). Likewise, individual effectors might interact preferentially with certain host protein families with respect to the rest of the network. This was tested using the same procedure.

### 3.5 Effector comparison

In order to compare the interaction patterns of different effectors, we computed all pairwise Jaccard similarities [Jaccard, 1912]. The Jaccard similarity is defined as the size of the intersection between sets (common interactions and non-interactions) divided by the size of the union of the sets (the whole set of interactions and non-interactions with host proteins that displayed at least one interaction in our Y2H assay). We also computed all possible pairwise sequence alignments (after removing signal peptides) using Clustal Omega v1.2.4 [Sievers et al., 2011] and calculated the corresponding sequence similarities. Sequence similarity is defined as the proportion of aligned residues with a log-odds score greater than 0 in the BLOSUM62 substitution matrix [Henikoff and Henikoff, 1992]. Finally, all dendrograms shown were constructed using average linkage hierarchical clustering with SciPy 1.3 [Virtanen et al., 2020] and (1 - similarity) as a distance metric.

### 3.6 PPI network comparison

We compared the host-pathogen interaction networks presented in [Mukhtar et al., 2011] and [Weßling et al., 2014] to ours. For many years, the Y2H system has been the gold standard for interactomics studies [Brückner et al., 2009]. Nevertheless, variation, as well as false negative and false positive interactions, can occur due to differences in the exact screening conditions, such as the yeast strain, vectors, incubation temperature and exact composition of selective media. For this reason, we focused on an overall comparison, rather than zooming in on unique, individual cases. Furthermore, the data of these previous studies contains no information about the host protein splicing variants used. Therefore, we had to collapse different splicing variants in our network into a single node. The only host protein within our PPI network with two different splicing variants was SEPALLATA4. One variant interacts with 5 different effectors, and the second interacts with a subset of 3 of these. Therefore, ignoring different splicing variants has a small or negligible effect in the comparison.

We selected the 753 host proteins that were present in all Y2H assays. Within this subset, 528 proteins did not interact with any effector from any species and were removed. Likewise, 7 effectors that did not show interactions with the remaining host proteins were removed. The final integrated network contains 118 effectors (19 from phytoplasma, 21 from *P. syringae*, 48 from *H. arabidopsidis* and 30 from *G. orontii*) and 225 host proteins. We performed a pairwise comparison of host protein degree per species using this network, taking into account host proteins that are relatively highly targeted by phytoplasma (≥4 interactions; this is the top 7% of host proteins targeted by phytoplasma) but not in other species. The same procedure was repeated using the larger subset of host proteins shared between our assay and the one performed on *G. orontii* effectors.

### 3.7 Phylogenetic tree construction

We constructed a phylogenetic tree of all TCP transcription factors in *A. thaliana* using their full length sequences with the ete toolkit (v 3.1.1) [Brückner et al., 2009, Huerta-Cepas et al., 2016]. The best model from JTT, WAG, VT, LG and mtREV was chosen using ProtTest [Abascal et al., 2005], and a maximum likelihood tree was built using PhyML [Guindon et al., 2010]. Reliability of branching was assessed with 100 bootstrap replicates.

For constructing the phytoplasma 16S rRNA and SAP11 trees, the 16S rRNA gene sequences and SAP11 protein sequences without the signal peptide were downloaded from NCBI, respectively (Table S2). Multiple sequence alignment (MSA) was performed with the MUSCLE algorithm using default parameters [Edgar, 2004] in MEGA v7 [Kumar et al., 2008]. The Maximum Likelihood algorithm was used for generating phylogenetic trees with the LG+I model and 1000 bootstrap replicates.. The resulting trees were formatted and annotated in FigTree v1.4.3 [Rambaut, 2012]. The SAP11 MSA was displayed using Boxshade [Hoffman and Baron, 1996].

### 3.8 Structural analysis of phytoplasma effectors

The structures of phytoplasma effectors were predicted using AlphaFold 2.2.0 [Jumper et al., 2021] using the monomer model and the reduced_dbs option. The top predicted models for each effector were used for subsequent analyses. Quality of the structures was assessed based on their median predicted local Distance Difference Test (pLDDT), a residue-wise confidence metric. The models are available in ModelArchive (modelarchive.org) with the accession code ma-saps.

Global structural similarities between different structures were evaluated with TM-align [Zhang and Skolnick, 2005]. We used a TM-score threshold of >0.5 to classify two structures as likely to be sharing the same fold [Xu and Zhang, 2010].

To identify local structural similarities between phytoplasma effectors, we used Geometricus, a structure embedding tool based on three-dimensional rotation invariant moments [Durairaj et al., 2020], which are discretized into so-called shapemers at a user-specified resolution. We followed the procedure of [Gordon et al., 2020] to uncover similarities between proteins that appear to share no common ancestry. The predicted structures were fragmented into shapemers both based on overlapping k-mers in the sequence (k = 20) and based on overlapping spheres surrounding each residue (radius = 15 °A). To ensure that similarities between structures, if any, would be significant, we used a high resolution of 7 to define the shapemers.

### 3.9 Generation of SAP06 transgenic plants

A synthetic plant codon-optimized ORF was created for AY-WB SAP06 by GenScript Biotech (New Jersey, US) and subcloned by Gateway-based cloning into the entry vector pDONR207 (Invitrogen, Thermo Fisher Scientific Inc., UK). The obtained donor vector was sequenced to confirm that the SAP06 ORF was correct and subsequently, a plant expression vector was generated by Gateway-based recombination. In this reaction, SAP06 was transferred into the destination vector pB7WG2 [Karimi et al., 2002], containing a CaMV 35S promoterdriven expression cassette. The obtained expression vector was transferred to Agrobacterium tumefaciens strain C58C1 and transformed into Arabidopsis Col-0 by the floral dip method [Karimi et al., 2002, Clough and Bent, 1998]. Primary transformants were obtained upon selective germination of seeds harvested from the transformed plants on agar plates containing 10 mg/l phosphinotrycin (PPT). From these individual lines, six 3:1 segregating lines were selected based on selective germination on PPT medium, followed by selection of homozygous lines in the next generation. qRT-PCR was used to select from these lines a transgenic line with a high ectopic expression level (SAP06#8) and a line with an intermediate expression level (SAP06#4). For all further phenotyping and molecular experiments, these two transgenic lines were used.

### 3.10 Expression analysis by qRT-PCR

Seedlings of the various transgenic SAP06 overexpression lines were grown under LD conditions (16h light, 8 h dark; 21 °C) on rockwool blocks. All above ground plant material from 7 day old seedlings was harvested, followed by RNA isolation using the InviTrap®Spin Universal RNA Mini Kit (Stratech, UK) according to the manufacturer’s instructions. The isolation was done in triplicate, with each sample consisting of plant material from at least five individual seedlings. DNA was removed by a DNAse treatment using the TURBO DNA-free^TM^ Kit (Invitrogen, Thermo Fisher Scientific Inc., UK). The iScript^TM^ cDNA Synthesis Kit (Bio-Rad, the Netherlands) was used to generate cDNA. qRT-PCR and calculation of relative expression was performed as described previously [Severing et al., 2018], but using SAND as reference gene. For the initial selection of SAP06 transgenic lines the primers PZN1326 ‘TGATGGTTGCTATCTCTAACACT’ and PZN1327 ‘GGCTTGTTCTGGTAGTTTCTTCT’ were used. For SAND the primers PDS2987 ‘TTCAAGAAGATGGAAGGTAATGATG’ and PDS2988 ‘CACCACT-CACTGATTTCCATTGCTTG’ were used; for the known DELLA targets *SPL3*, *EXP8*, *PRE5*, and *PRE1* the primers that were described previously [Park et al., 2013]. Statistical significance of observed differences in DELLA target gene expression was evaluated by performing a 2-way ANOVA (using both genes and lines as factors) followed by Tukey’s honest significance test in statsmodels v0.13.5 [Seabold and Perktold, 2010].

### 3.11 Phenotyping of SAP06 ectopic expression lines

For the phenotyping of seed dormancy, seeds of the two selected SAP06 ectopic expression lines and of wild type Col-0 were sown on blue filter paper wetted with MQ water in 10 cm Petri dishes, followed by the scoring of the percentage of germination after 6 days of incubation at either 21 °C or 25 °C. Each measurement is based on scoring germination for at least 66 seeds. The remaining seeds were stored at room temperature (20-21 °C) and sown in a similar way after one and two weeks of storage. Germination of these batches was scored as well at 6 days after the start of the germination assay. Since all seeds were germinating after two weeks of storage, we stopped scoring for dormancy at this point. To assess whether the differences between lines are significant, we conducted logistic regression using statsmodels v0.13.5 [Seabold and Perktold, 2010]. We included as variables the line to which each seed belongs (encoded as two dummy variables), the temperature at which the seed was incubated, and the number of weeks of storage after seed harvest & before sowing For the characterization of potential effects on vegetative development, seeds of the two SAP06 ectopic expression lines and of Col-0 wild type were sown on wetted filter paper and stratified for three days at 4 °C. Subsequently, seeds were sown on Rockwool plugs. Twenty seedlings of each genetic background were grown under long day conditions (16/8, light/dark) at 21 °C. Three, four and five weeks after germination, pictures were taken of representative individual plants of each genetic background.

## 4 Results & discussion

### 4.1 Widespread interactions between phytoplasma effectors and *A. thaliana* transcription factors

To obtain a better understanding of the molecular mechanisms through which phytoplasmas infect and manipulate their host, we performed a large-scale matrix-based yeast two-hybrid (Y2H) screening to find putative interaction partners of 21 phytoplasma effectors in an existing library containing 1980 *A. thaliana* transcription factors and transcriptional regulators [Pruneda-Paz et al., 2014]. The library contains a broad and diverse set of host proteins, which provide comprehensive coverage of *A. thaliana* transcription-related proteins. Proteins in the library belong to 95 different families, with no clear predominating group (Fig. S1). Although *in planta* activity has not been validated for all 21 studied and putative effector proteins, such activity has been suggested based on various criteria [Bai et al., 2009, MacLean et al., 2011]. For this reason, and for simplicity, we will hereafter refer to these 21 putative effectors as effectors. Seventeen of the effector proteins were chosen from phytoplasma strain AY-WB based on significant up-regulation of the genes encoding these effectors in the phytoplasma infecting *A. thaliana*, compared to expression levels of these phytoplasma genes in the vector *Macrosteles quadrilineatus* [MacLean et al., 2011]. The other four selected SAP proteins represent orthologs of the previously characterized AY-WB SAP11 and SAP54 proteins from other phytoplasma isolates. Initially, an auto-activation test was performed for the 21 effector protein used as bait proteins. Autoactivation was only found for SAP06 and SAP48 on selective SD medium supplemented with up to 15 mM 3-amino-1,2,4-triazole (3-AT). For this reason, these baits were screened in the Y2H assay with the more selective Adenine reporter and on medium supplemented with higher concentrations of 3AT. For the previously studied AY-WB SAP11 and SAP54 proteins, we confirmed various previously found interactions, for example between SAP11 and TCP2, TCP7 and TCP13 [Sugio et al., 2011a, 2014]. Likewise, interactions were identified between SAP54 and different MIKC MADS-box proteins involved in floral organ specification and determination of flowering time (such as AP1, SEP3 and SOC1, respectively) [MacLean et al., 2014] and between SAP05 and different SBP and GATA transcription factors [Huang et al., 2021]. Note however that some interactions were not confirmed in our matrix-based screen (e.g. the interaction between SAP54 and the MADS-box protein AGAMOUS-LIKE 12). The most likely reason for this is the different setup and more stringent conditions to select for interactions in this study, which may lead to false negatives.

For all tested effector proteins, multiple new interacting host proteins were identified, and the resulting pathogen-host PPI network represents 979 interactions, involving 536 (28%) of the screened host proteins (Fig. 1a; Table S1). The identified interactions have been submitted to the IMEx consortium (http://www.imexconsortium.org) through IntAct [Orchard et al., 2014] and assigned the identifier IM-28211. All of the effectors displayed at least two interactions, but the distribution of degrees (i.e., the number of interactions for each protein) is broad (Fig. 1b). Remarkably, two related effectors, SAP48 and SAP06, interacted with a large number of host proteins (250 and 242, respectively) from multiple protein families. Many of these host proteins did not interact with any other effector we tested, revealing the specificity of the different SAP proteins and that the assay can distinguish differences in interactions. However, given that both SAP06 and SAP48 exhibited autoactivation in our initial screening, interactions with these two effectors should be interpreted carefully despite the stringent conditions used. Nonetheless, given that SAP06 and SAP48 are phylogenetically related (42.4% sequence similarity) but have highly dissimilar interaction patterns, it would be unexpected that all these interactions represent false positives and could be solely attributed to the observed autoactivation.

**Figure 1:**
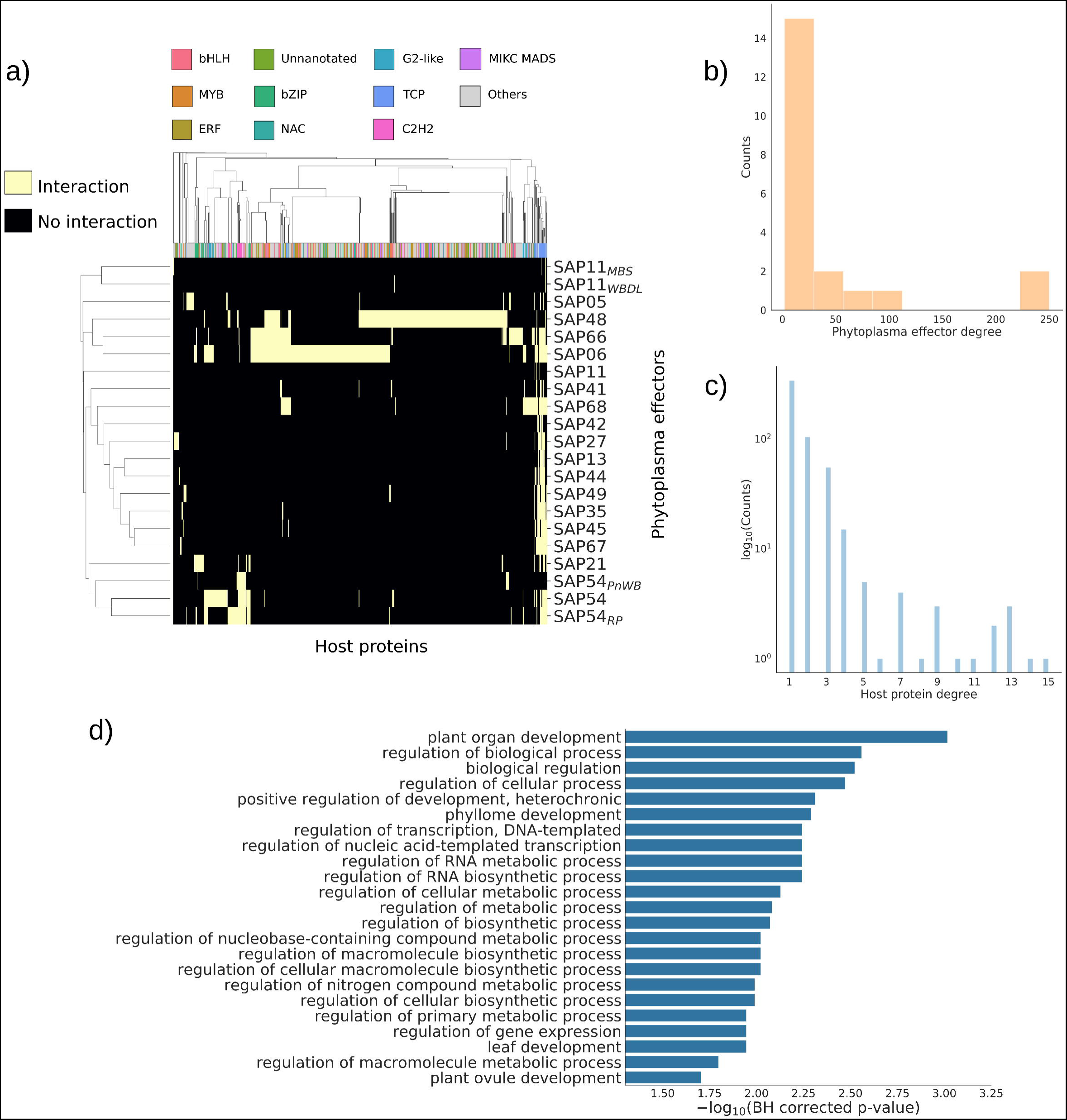
Overview of the obtained phytoplasma-*Arabidopsis* interaction network. (a) Clus-termap of the interactions, grouping host proteins (horizontal) and effectors (vertical) with similar interaction patterns. Colors indicate family membership of the corresponding host proteins for the 10 most abundant host protein families in the network. Clustering was performed using Jaccard distance as a metric. (b) Histogram of candidate effector degrees (the number of interactions detected for each protein). (c) Histogram of host protein degrees (with counts in log scale). (d) Significantly enriched biological process GO terms in the interaction network, sorted from lowest (above) to highest (below) p-value. Note that the horizontal axis starts at the statistical significance threshold.

Additionally, large variation was found in the number of interactions per host protein. Although 63% of the interacting host proteins only bind one effector, some of them are clearly highly targeted (Fig. 1c). Remarkably, many of these highly-targeted host proteins belong to the TCP transcription factor family (Fig. 1a), and the 10 host proteins with the highest degree are all TCPs. Other host proteins with a high degree include LBD15, a protein involved in shoot apical meristem development [Sun et al., 2013], and two proteins of unknown function: a homeobox and a NAC transcription factor (AT4G03250.1 and AT3G12910.1, respectively). This high number of interactions suggests important roles of these transcription factors in phytoplasma infection.

We performed GO enrichment analysis to determine whether there is an overrepresentation of host proteins in the network that participate in particular functions with respect to the library. Aside from general, high-level terms, this revealed a highly significant enrichment in different biological process terms related to plant development (Fig. 1d), and in particular in terms related to the development of phyllomes (i.e., organs homologous to leaves or derived from leaves, such as flowers), which is consistent with known phytoplasma infection symptoms. Altogether, this indicates that the network captures biologically relevant interactions.

### 4.2 Multiple unrelated effectors specifically target TCP transcription factors

To investigate whether phytoplasma effectors interact specifically with certain protein families, we evaluated whether particular families are enriched in the network with respect to the library. We found that only one family, TCP transcription factors, is significantly enriched in the network (adjusted p = 2.5 × 10^−2^). Of the 24 TCPs present in the *A. thaliana* genome, which are all present in the library, 20 are found in the network. Furthermore, these interactions are distributed over the whole TCP family, without particular enrichment of a specific class or subclass (Fig. 2a). TCPs form an ancient, plant-specific family of transcription factors, found from green algae to eudicots [Floyd and Bowman, 2007, Navaud et al., 2007]. Although TCPs were initially linked to plant growth and development [Martín-Trillo and Cubas, 2010], research in recent years has uncovered that they also participate in the plant immune response [Lopez et al., 2015, Li, 2015]. These properties seem to make them important targets for pathogens. Some TCPs have previously been found to be targeted by the phytoplasma effector SAP11 [Sugio et al., 2011a, 2014, Chang et al., 2018, Wang et al., 2018, Pecher et al., 2019], and by effectors from evolutionarily distant pathogens [Mukhtar et al., 2011, Weßling et al., 2014].

**Figure 2:**
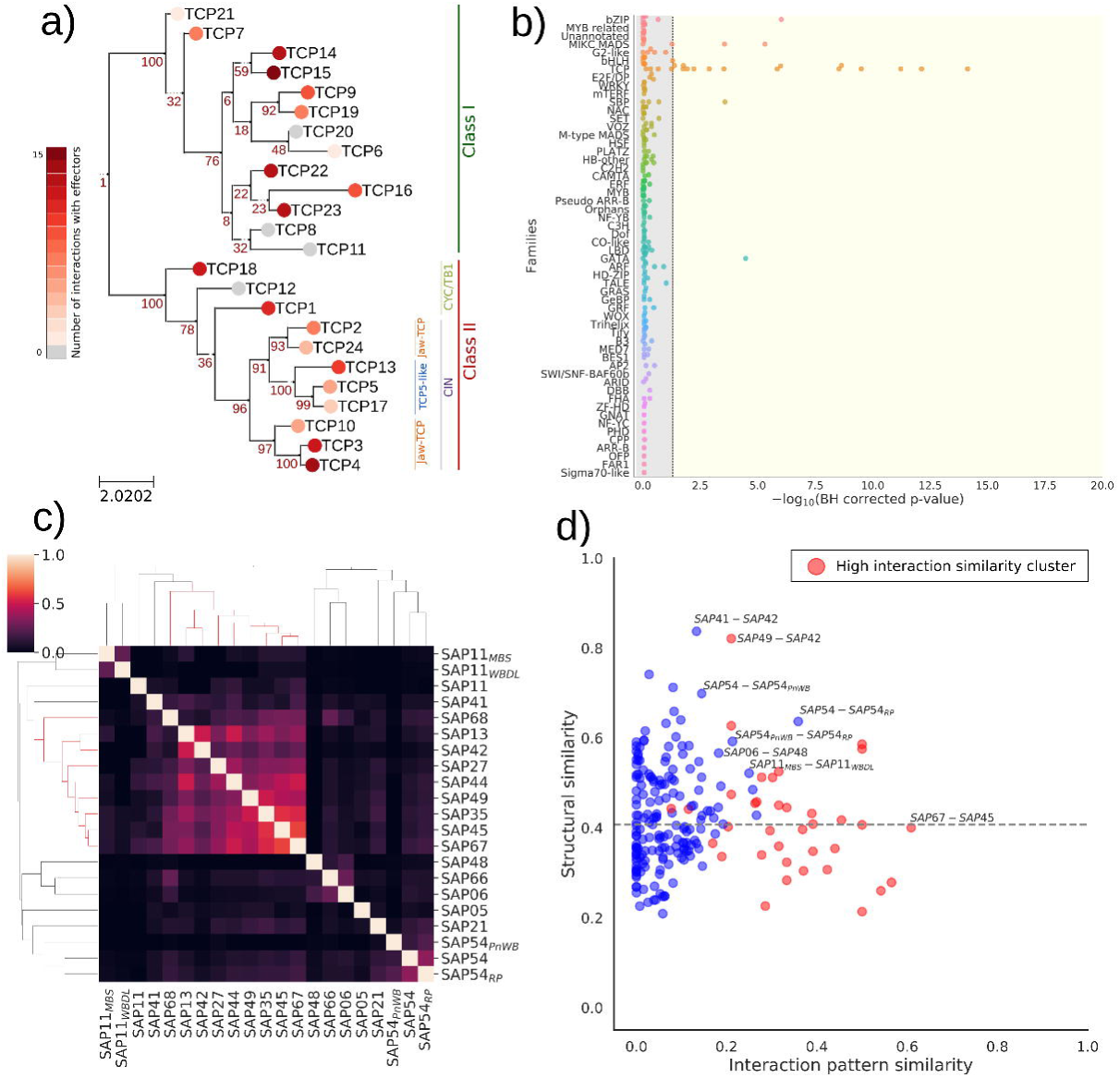
Zooming into specific host-pathogen interactions. (a) Phylogenetic tree of all fulllength TCP factors in *A. thaliana*. Leaf node color indicates the number of interactions with phytoplasma effectors. Numbers at internal nodes indicate bootstrap branching support. Class assignment according to [González-Grandío and Cubas, 2016]. (b) Overview of overrepresentation in interactions for different host protein families across individual effectors (indicated by each dot). The vertical dashed line indicates the statistical significance threshold (adjusted p <0.05). (c) Clustermap of effector interaction similarities, measured by Jaccard distance. The cluster of TCP-specific effectors with high interaction similarities is highlighted in red in the dendrogram. (d) Relationship between effector interaction similarity and structural similarity (quantified as TM-scores). Effectors belonging to the cluster of sequences with similar interaction patterns are marked in red; outliers and interesting examples are annotated with the pair they represent. The horizontal dashed line indicates median structural similarity.

To obtain a more precise picture, we tested whether individual effectors interact predominantly with specific host protein families. Strikingly, we find that 15 effectors (71%) are enriched in interactions with TCPs (Fig. 2b). This set includes SAP54 and its ortholog SAP54*_RP_*, whose interactions with TCPs had previously not been identified. The fact that there is a high level of functional redundancy within the TCPs [Sugio et al., 2011a, Danisman et al., 2013], together with their interactions with numerous phytoplasma effectors, suggests that suppression of TCP activity is crucial for phytoplasmas, and that it can only be achieved effectively by targeting many TCPs with multiple effectors.

However, some effectors display different specificity: SAP05 has an overrepresentation of interactions with SBPs and GATAs, confirming other studies of this effector [Huang et al., 2021]. Furthermore, its set of interactors is enriched in proteins (all SBPs) involved in anther development (adjusted p = 8.2 × 10^−3^). AY-WB infected *Arabidopsis* plants have abnormal anthers that do not produce pollen (MacLean et al., 2011). This suggests that SAP05 is involved in making the infected host sterile. Additionally, one of the TCP-enriched effectors, SAP21, is enriched in interactions with bZIPs. Finally, as expected given their previously described interactions [MacLean et al., 2014], the SAP54s are enriched in interactions with MIKC MADS-box proteins.

These analyses revealed broad general patterns shared between different effectors, which led us to evaluate how redundant their interaction patterns are. Overall, they tend to be very different from each other: the median similarity is 0.06, where 0 means no overlap in interactions and non-interactions and 1 complete overlap. Grouping effectors by interaction similarity reveals, aside from the expected clusters of orthologous effectors from different strains, a clear cluster of nine effectors with similar interaction patterns (Fig. 2c, highlighted in red in the dendrogram), all enriched in interactions with TCPs.

To understand whether these similarities in interaction patterns might emerge from structural similarities or homology, we used AlphaFold2 [Jumper et al., 2021] to predict the structures of all effectors in our library, which yielded structures with overall high confidence (80.95% of structures have a median pLDDT ≥ 75) (Fig. S2). We also compared the recently experimentally solved structure of SAP05 [Liu et al., 2023] and its predicted counterpart, showing a high agreement (root mean square deviation = 0.65) (Fig. S2d). Global comparison of the structures shows that most structures do not share the same fold (75% TM-score ≤ 0.5, ignoring comparisons between SAP11 orthologs and between SAP54 orthologs [Zhang and Skolnick, 2005, Xu and Zhang, 2010]) (Fig. S3a). This trend of general dissimilarity is found in both the high interaction similarity cluster (from now on referred to as HISC) and the rest of the effectors (Fig. S3b). Interestingly, despite having similar interaction patterns, these effectors tend to have both low structure and sequence similarities (Fig. 2d, Fig. S4), suggesting that they are, overall, phylogenetically unrelated.

Although globally the effector structures may be dissimilar, there may be common structural or sequence elements to allow them to bind TCPs. We created embeddings of the structures using Geometricus [Durairaj et al., 2020], an algorithm that creates representations of protein structures based on overlapping sequence k-mers and overlapping spheres. These are aggregated into shapemers, which represent a set of similar structural fragments. We find that the effectors do not clearly cluster into the HISC and the background according to the shapemers (Fig. S3c). Furthermore, there is a large variety of shapemers exclusive to each group, as expected from the overall dissimilarity between structures. Inspecting the shapemers exclusive to the HISC shows no clear commonalities (Fig. S3d). Finally, the frequencies of shapemers that are common between the two groups do not show a clear substructure that could explain any common binding mechanism. At most, we can identify 2 different radius-based shapemers found in 4 out of 9 effectors in the HISC and in none of the other effectors. An analogous sequence-based approach using discriminant motif analysis [Bailey, 2011] on the clustered sequences, using effectors with no specificity for TCPs as a control, yields no statistically significant results. From these results, we conclude that it is very unlikely that there is a similar binding site across effectors in the HISC. Overall, this indicates a certain degree of functional redundancy between these effectors.

According to the Red Queen hypothesis, mutually antagonistic interactions between host and pathogen are predicted to lead to evolutionary arms races [Van Valen, 1973]. It is known that many pathogens produce host-mimetic molecules in order to exploit it [Bailey, 2011, Elde and Malik, 2009, Via et al., 2015], and it has been postulated that phytoplasma effectors mimic the structures of their targets in order to interact with them [Rümpler et al., 2015]. Conversely, host proteins involved in pathogen interactions tend to be under adaptive selection [Mukhtar et al., 2011, Weßling et al., 2014, Slodkowicz and Goldman, 2020]. Nonetheless, given the large number of unrelated effectors targeting them, it could be diffcult for targeted TCP host proteins to evolve in a way that prevents interactions with effectors. Changes at the interaction interface that weaken interactions with an effector might also weaken physiologically important interactions. Additionally, multiple unrelated effectors with similar interaction patterns could still be able to interact with the host protein, rendering such adaptive changes moot. In this way, phytoplasmas could mount a very effective attack that hampers the evolution of a defense by setting an evolutionary trap in sequence space.

Interestingly, orthologous effectors from different strains have variable interaction similarities. We observe that interactions with MIKC MADS-box proteins responsible for floral organ identity, such as AP1 or SEP3, are conserved across different SAP54s. In contrast, interactions with MIKCs involved in flowering time are less conserved, except for interactions with SOC1 (Fig. S5a). Floral organ specification mechanisms are strongly conserved, while flowering time is linked to adaptation of a plant to its environment. Furthermore, while interactions of SAP54 and SAP54*_RP_* with TCPs are highly conserved (Fig. S5b), no such interactions are detected for the SAP54*_P nW B_* ortholog. Altogether, this strongly suggests coevolution with orthologous host proteins, eventually leading to divergence at the interaction interface.

### 4.3 Phytoplasma SAP11 homologs show binding specificity towards TCP subclasses

Phytoplasma SAP11 effectors show interactions with specific TCP proteins in our screen, and were previously found to interact with plant TCPs as well [Sugio et al., 2011a, 2014, Chang et al., 2018, Wang et al., 2018, Pecher et al., 2019]. Plant TCP transcription factors group into several (sub)classes that antagonistically modulate plant growth and development by competitively binding similar cis-regulatory modules called site II elements. Phytoplasma SAP11 effectors were found to show binding specificities towards TCP (sub)classes, e.g. SAP11*_AY W B_* interacts with class II TB1 and CIN/jaw-TCPs, whereas SAP11*_MBS_* interacts with only TB1 TCPs (Table S1), and neither interact with class I TCPs [Sugio et al., 2011a, 2014, Pecher et al., 2019]. Here, we identified more SAP11 homologs from publicly available sequence data of diverse phytoplasmas. The alignment of these SAP11 homologs showed differences in the protein region known to interact with TCPs (Fig. 3a). To further investigate phytoplasma effector interaction specificity with TCPs and possible differences in evolutionary trajectories, we cloned additional SAP11 effectors from diverse phytoplas-mas (Tables S2 & S3) and tested their binding specificities to a selected subset of *A. thaliana* class I and II TCPs by Y2H.

**Figure 3:**
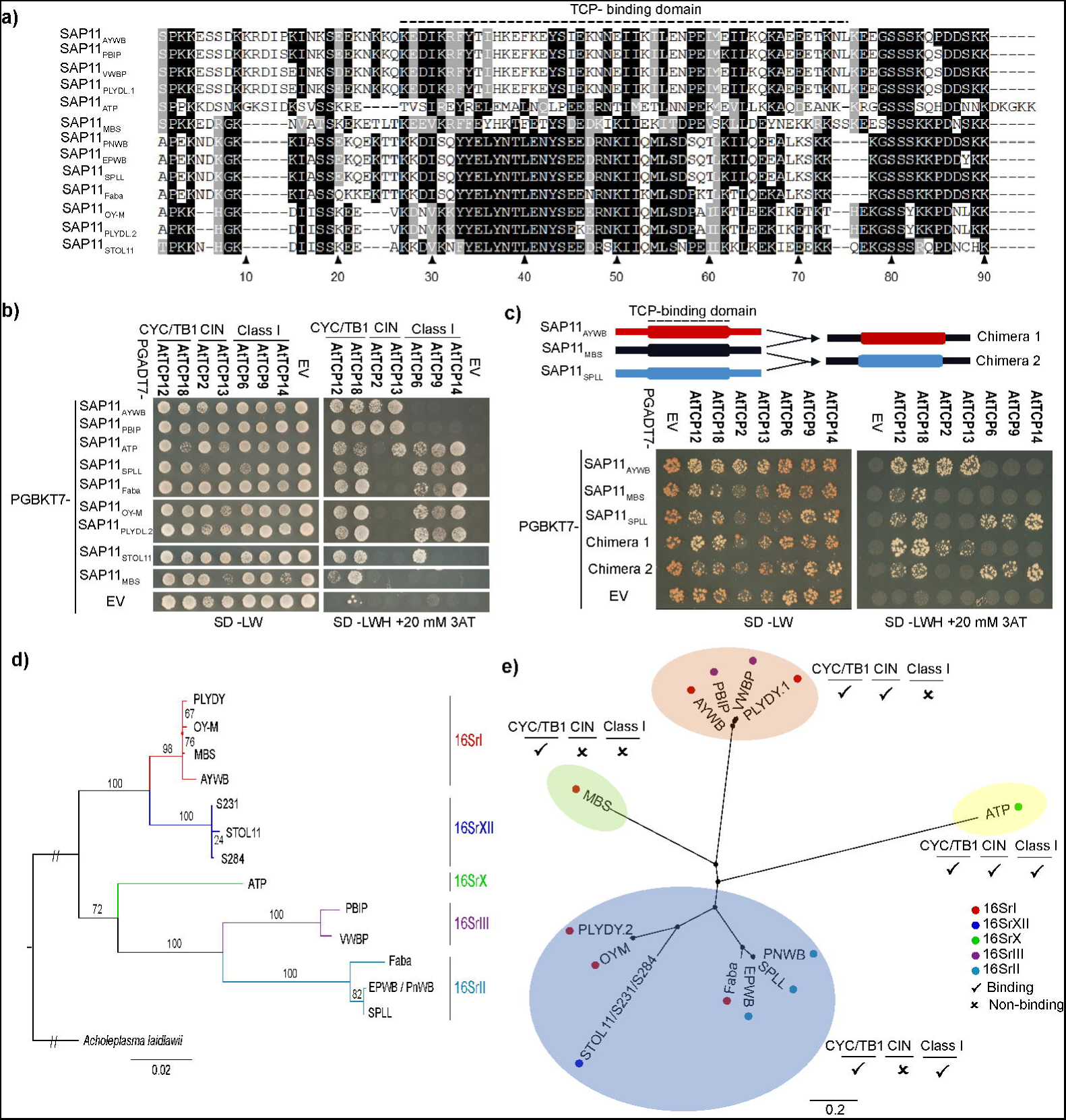
SAP11 homologs show distinct interaction specificities for CIN and class I TCPs. (a) Alignment of SAP11 homologs. The dashed line indicates the TCP-binding domain on SAP11*_AY W B_*. (b) Yeast two-hybrid analysis of the interaction of phytoplasma SAP11 homologs and representative members of the three TCP subclasses in *Arabidopsis thaliana*. SD-LW, yeast synthetic drop-out without leucine and threonine. SD-LWH, yeast synthetic drop-out without leucine, threonine and histidine. 3AT, 3-Amino-1,2,4-Triazol. EV, empty vector. (c) SAP11-binding specificity to CIN versus class I TCPs is determined by a 50 residue region. Upper panel, schematic illustration of SAP11 homologs or SAP11 chimeras. Lower panel, Y2H analysis. (d) Known phytoplasmas that contain SAP11 effector genes belong to five distinct 16Sr subgroups. Phytoplasmas clades are color coded according to their subgroups. (e) Phylogeny of SAP11 homologs combined with their binding specificities towards the three TCP subclasses. Numbers on the internal branches indicate the level of bootstrap support (percentage).

This assay confirmed that SAP11*_AY W B_* and SAP11*_MBS_* interacted with specific class II TCPs, including TB1 and CIN/jaw-TCPs. SAP11 homologs SAP11*_P LY DY._*_1_, SAP11*_V W BP_* and SAP11*_BP IP_* had similar binding specificity to class II TCPs as SAP11*_AY W B_* did (Fig. 3b). Surprisingly, we discovered that some SAP11 homologs (SAP11*_P nW B_*, SAP11*_SP LL_*, SAP11*_EP W B_*, SAP11*_F aba_*, SAP11*_OY_* _−_*_M_*, SAP11*_P LY DY._*_2_ and SAP11*_ST OL_*_11_*_/S_*_231_*_/S_*_284_) interacted with class I TCPs but not with class II CIN-TCPs, whereas SAP11 homolog SAP11*_AT P_* interacted with members of all three TCP (sub)classes (Fig. 3b). Nevertheless, all tested SAP11s share an interaction with class II TB1 TCPs (Fig. 3b). Moreover, in the phylogenetic analyses of SAP11 homologs, the SAP11 proteins cluster based on their interaction specificities with TCPs, rather than based on the phytoplasma 16S rRNA phylogeny (Fig. 3d, e). To independently provide evidence that the distinct sequences among the SAP11 homologs (Fig. 3a) are involved in TCP binding, we generated SAP11 chimeras using the SAP11*_MBS_* sequence as a backbone (Fig. 3c). This showed that the region designated as TCP-binding (Fig. 3a) is indeed involved in determining the specificity of SAP11 binding to TCP (sub)classes. Taken together, these data show that SAP11 effectors have evolved binding specificity for specific (sub)classes of TCP transcription factors. Therefore, despite the fact that TCPs were identified as interactors of many SAPs, SAP-TCP interactions do show specificity.

### 4.4 Analysis of interaction partners predicts effector-induced phenotypes

In order to study whether interactions with a defined set of host proteins might be used to predict potential functional consequences of the effectors, we tested for overrepresentation of particular GO terms in the effector-specific sets of host interactors (Table S4). As already indicated, terms related to plant development are highly overrepresented, and for seven of the SAPs the top ranking GO term is associated with development. However, specificity can also be found at this level: for example, defence response is the top-ranking term for SAP45 and SAP49. For the highly reactive SAP06 protein, the analysis of the interactors revealed enrichment in interactions with multiple proteins from different families involved in leaf growth and regulation of seed dormancy. Related to the latter biological process, the presence of multiple DELLA proteins, which are well-known repressors of gibberellic acid-induced seed germination [Ravindran and Kumar, 2019], stands out.

Given that SAP06 displayed autoactivation in our initial screenings, we first set out to confirm the interactions with DELLA proteins. We used BiFC in *Nicotiana benthamiana*, a methodology orthogonal to Y2H, to test interactions with four different DELLA proteins. The experiment confirmed all four interactions (Fig. S6).

Inspired by these findings and having confirmed the interactions with DELLA proteins, we tested empirically whether SAP06 has an effect on plant growth and seed dormancy in line with the expectations based on the outcomes of overrepresentation analysis. We created transgenic *A. thaliana* plants with ectopic expression of SAP06 and selected two lines with different levels of SAP06 expression (Fig. S7). Phenotyping of these transgenic lines during the vegetative stage of development indeed revealed stunted growth compared to Col-0 control plants, with an effect proportional to the level of SAP06 expression (Fig. 4a; Fig. S7). Subsequently, we also tested whether SAP06 affects the expression of four known DELLA target genes [Park et al., 2013]. We found that the expression of one of these, *PRE1*, is significantly downregulated in the line with the highest SAP06 expression (Fig. S8). Although the expression changes in the other three measured genes do not meet the criteria for statistical significance, DELLA target genes generally seem to be downregulated in the line with the highest SAP06 expression. Performing further phenotyping analyses, we found statistically significant altered levels of seed dormancy as well in the transgenic line with high ectopic SAP06 expression (Fig. 4b; p-value of SAP06#8 model coefficient = 3.7 × 10^−10^; log-likelihood ratio p-value of the logistic regression = 1.2 × 10^−18^). All together, the different observations of stunted vegetative growth, increased seed dormancy, and significant downregulation of *PRE1*, consistently point to stabilization of the DELLA proteins upon interaction with SAP06. An effect on seed dormancy due to a phytoplasma infection has been previously noted in the form of vivipary in tomato [Wei et al., 2019]. However, in that case less dormancy and premature germination was observed, while here in *Arabidopsis* we see more seed dormancy as a result of the ectopic SAP06 expression. Note, however, that this is a phenotype upon artificial ectopic expression of a single effector protein in comparison to a phytoplasma infection in the previous study. Hence, our results reveal a potential function of the SAP06 effector protein and it is very well possible that SAP06 interacts with DELLA proteins to interfere with other functions, such as immunity, and the balance and trade-off between immunity and growth [Davière and Achard, 2016]. Nevertheless, these findings highlight how the PPI network can be interpreted to generate insights into the function of individual effectors.

**Figure 4:**
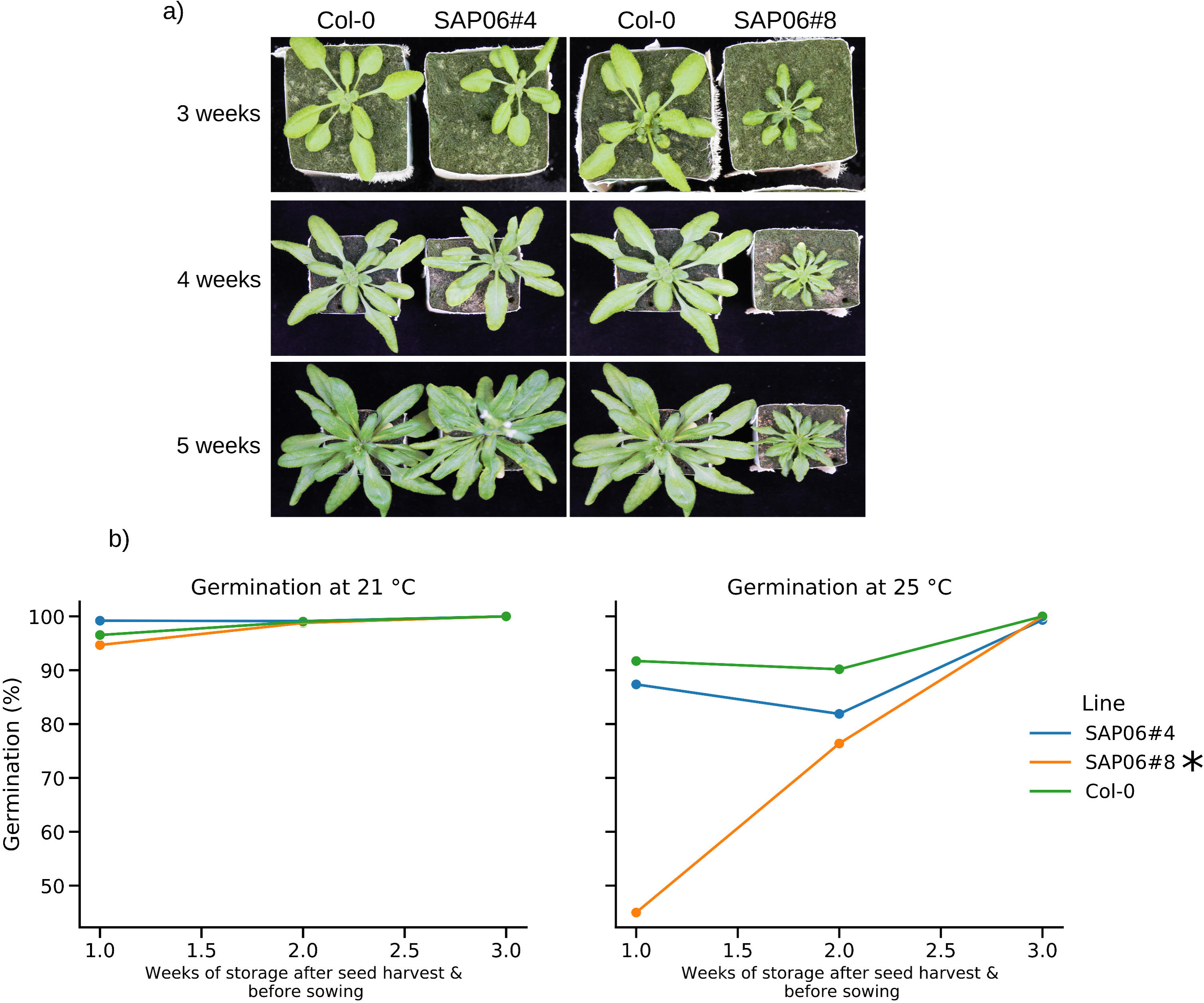
Phenotypic alterations upon ectopic expression of AY-WB SAP06 in *A. thaliana* Col-0 background. (a) Effect of ectopic SAP06 expression on growth and development in the vegetative stage, three, four and five weeks after germination. To the left, comparison between Col-0 and a line with a medium level of SAP06 expression; to the right, comparison between Col-0 and a line with a high level of SAP06 expression. (b) Effect of ectopic SAP06 expression on seed dormancy in the same transformed lines. Germination percentage after incubation at 21 degrees C (left) and 25 degrees C (right) upon direct sowing or after storage of seeds at room temperature for one or two weeks. * next to line name indicates statistically significant differences.

### 4.5 Phytoplasma has developed specific strategies to target host development

Prior research has determined PPI networks between *A. thaliana* proteins and the effectors of different, evolutionarily distant pathogens: the bacterium *Pseudomonas syringae*, the fungus *Golovinomyces orontii* and the oomycete *Hyaloperonospora arabidopsidis* [Mukhtar et al., 2011, Weßling et al., 2014]. These studies found that effectors produced by pathogens from different kingdoms target overlapping sets of plant proteins, indicating the existence of a conserved host-pathogen interface. This includes proteins involved in defense response, but also e.g. auxin and salicylic acid signaling. In order to determine whether phytoplasma fits this picture, we compared our PPI network to the networks determined by [Mukhtar et al., 2011, Weßling et al., 2014] using the common subset of host proteins screened in all assays (Fig. S9a). As expected, the subset of host proteins that interacts with effectors from all pathogens (including phytoplasma) is enriched in proteins involved in stress and defense response (Fig. S9b). Remarkably, a high number of interactions with TCP14, which promotes disease resistance [Yang et al., 2017], is conserved across all pathogens. However, clustering of all effectors according to their interaction patterns separates most phytoplasma effectors from the others (Fig. 5a). Clearly, phytoplasma effectors tend to have far more interactions with this subset of proteins than other species’ effectors (Fig. 5a,b). This indicates that phytoplasma effectors diverge in their interaction patterns from effectors secreted by other pathogens.

**Figure 5:**
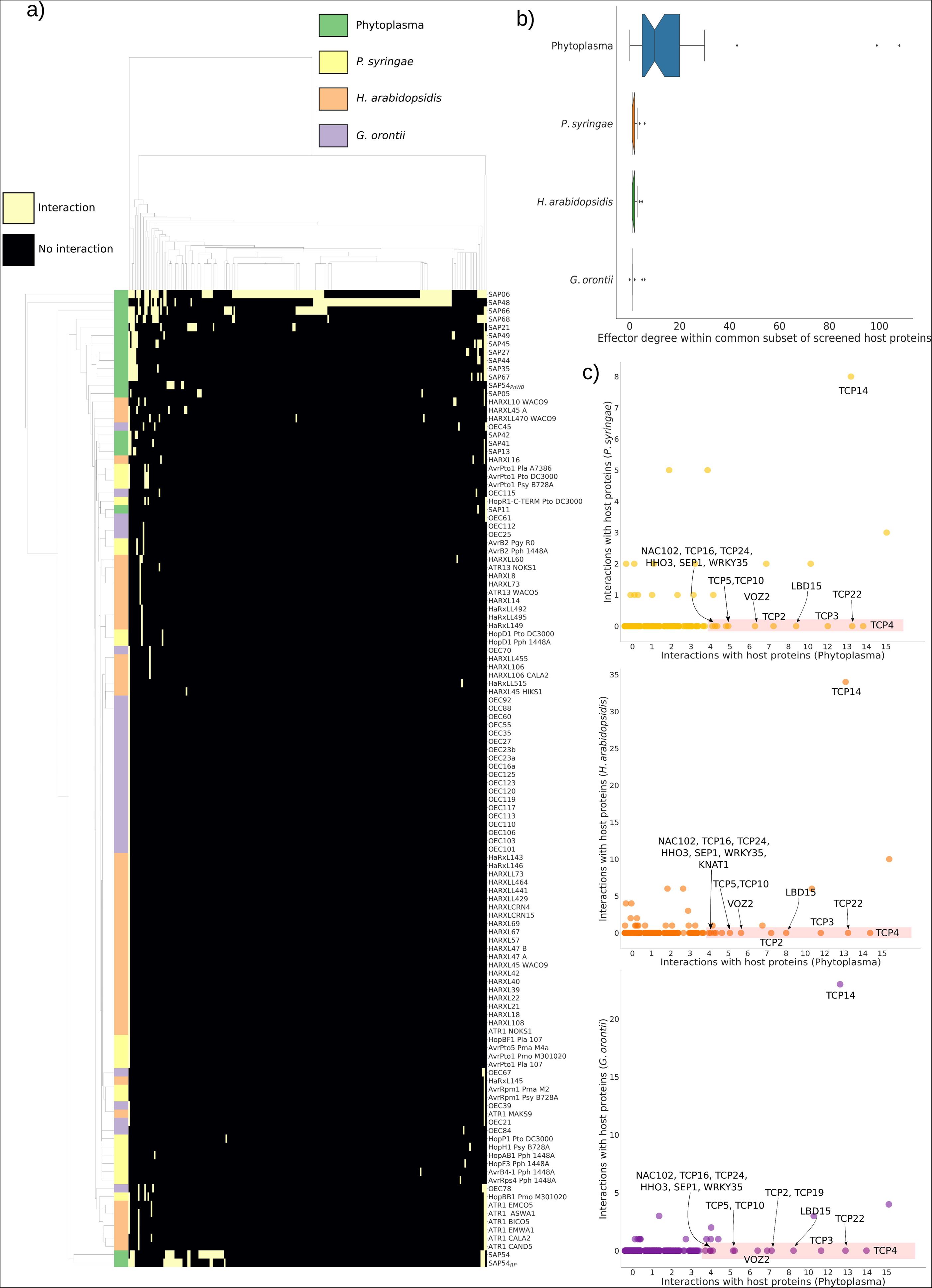
(a) Clustermap of interaction patterns in different host-pathogen PPI networks within the subset of screened host proteins shared by all assays. Clustering was performed using Jaccard similarity as a metric. (b) Boxplot of the number of interactions of effectors within the common host protein subset. Notches indicate the width of a 95% confidence interval of the median degree, determined by 1000 bootstrap resamples. (c) Comparison of host protein degrees between phytoplasma and *P. syringae* (left), *H. arabidopsidis* (center) and *G. orontii* (right). Host proteins highly targeted by phytoplasma but not by individual pathogens are highlighted. Also TCP14 is highlighted, which appeared to be a target of effectors of all investigated pathogens.

Due to possible technical variability caused by differences in the exact screening conditions, we focused our network comparison on host proteins that are highly targeted by phytoplasma effectors but did not interact with the effectors of other, individual pathogens. We find that there is little variation in this set of proteins across pathogens (Fig. 5c). This set contains a core of 12 proteins that remains constant. This highly targeted core is enriched in many terms related to regulation of development and phyllome development (Fig. S9b). The core includes multiple TCPs (including all of the CIN/jaw-TCPs) and other development-related proteins from different families, such as SEP1 and LBD15. It also includes the flowering time regulator VOZ2 [Yasui et al., 2012]. We repeated the analysis using the larger subset of host proteins shared between our assay and the one performed with *G. orontii* effectors, reaching the same conclusions (Fig. S10). Although previous studies have found a certain level of enrichment in targeting development-related proteins in *H. arabidopsidis*, the intense focus on targeting development-related proteins in phytoplasmas is unprecedented. Furthermore, we find only one protein within this specific subset of *A. thaliana* proteins (TCP20, Fig. S10c) that does not interact with a phytoplasma effector but interacts with effectors of a different pathogen. Put together, these observations indicate that phytoplasma has developed idiosyncratic molecular mechanisms that specifically target development in order to alter its host.

## 5 Conclusions

We have experimentally determined interactions between phytoplasma effectors and *A. thaliana* transcription factors and transcriptional regulators. Many of the assayed effectors have not been previously studied, and this work presents insights into their putative molecular mode of action. We find that interactions with host transcription factors enriched in development regulation are pervasive amongst phytoplasma effectors. Furthermore, compared to other phytopathogens, phytoplasmas appear to have unique targets. The determined network can be used to predict functional consequences of specific effectors, and our interaction data and findings can be further utilized to formulate mechanistic hypotheses regarding individual effectors. Future molecular data on phytoplasma, such as proteomics quantification of protein abundance upon infection, can be integrated with the presented PPI network and predicted effector structures to advance our systems-level view of phytoplasma pathogenesis and yield further insights.

Remarkably, our analysis shows that many effectors interact with TCP transcription factors, almost all of which are targeted, but with specificity. This indicates that suppressing the activity of TCPs, which resemble a crossroad between development and immunity, is crucial for the infection strategy of phytoplasma. Furthermore, while many of these TCP-specific effectors have similar interaction patterns, they bear little sequence similarity to each other. This functional redundancy between very different sequences could make it difficult for the affected host proteins to evolve to avoid these interactions, since they are targeted by multiple seemingly unrelated effectors, while they have to maintain their physiological interactions.

Phytoplasmas have coevolved with seed plants and their insect sap-feeding vectors [Cao et al., 2020], depending on both these hosts for spreading, and likely have optimized strategies to modulate both organisms and their interactions. This study serves as a stepping stone for a more global understanding of phytoplasmas and how they interact with and manipulate their hosts.

## Acknowledgments

The project was supported by the research school Experimental Plant Sciences (EPS), Wageningen, the Netherlands, (grant EPS3 4a 124), the Human Frontier Science Program grant RGP0024/2015 (to S.A.H. and R.G.H.I.), the Biotechnology and Biological Sciences Research Council (BBSRC) Institute Strategic Programme Grant BBS/E/J/000PR9797 (to JIC) and BBSRC PhD student fellowship of S.C. with additional support from the John Innes Foundation and the Gatsby Charitable Foundation.

We are grateful to Dr. Amalia Diaz Granados for advice on the BiFC assay and for providing the positive control for this experiment. We also wish to thank Dr. Jü rgen Jänes for running the AlphaFold2 prediction pipeline on our behalf.

## References

Federico Abascal, Rafael Zardoya, and David Posada. ProtTest: selection of best-fit models of protein evolution. Bioinformatics, 21(9):2104–2105, May 2005.

Xiaodong Bai, Valdir R Correa, Tania Y Toruno, El-Desouky Ammar, Sophien Kamoun, and Saskia A Hogenhout. AY-WB phytoplasma secretes a protein that targets plant cell nuclei. Mol. Plant. Microbe. Interact., 22(1):18–30, January 2009.

Timothy L Bailey. DREME: motif discovery in transcription factor ChIP-seq data. Bioinformatics, 27(12):1653–1659, June 2011.

Yoav Benjamini and Yosef Hochberg. Controlling the false discovery rate: A practical and powerful approach to multiple testing, 1995.

Assunta Bertaccini. Phytoplasmas: diversity, taxonomy, and epidemiology. Front. Biosci., 12:673–689, January 2007.

Anna Bruckner, Cecile Polge, Nicolas Lentze, Daniel Auerbach, and Uwe Schlattner. Yeast two-hybrid, a powerful tool for systems biology. Int. J. Mol. Sci., 10(6):2763–2788, June 2009.

Yanghui Cao, Valeria Trivellone, and Christopher H Dietrich. A timetree for phytoplasmas (mollicutes) with new insights on patterns of evolution and diversification. Mol. Phylogenet. Evol., 149:106826, August 2020.

Shu Heng Chang, Choon Meng Tan, Chih-Tang Wu, Tzu-Hsiang Lin, Shin-Ying Jiang, RenCi Liu, Ming-Chen Tsai, Li-Wen Su, and Jun-Yi Yang. Alterations of plant architecture and phase transition by the phytoplasma virulence factor SAP11. J. Exp. Bot., 69(22): 5389–5401, November 2018.

S J Clough and A F Bent. Floral dip: a simplified method for agrobacterium-mediated transformation of arabidopsis thaliana. Plant J., 16(6):735–743, December 1998.

N E Cortés-Martínez, E Valadez-Moctezuma, L X Zelaya-Molina, and N Marban-Mendoza. First report of ‘candidatus phytoplasma asteris’-related strains infecting lily in mexico. Plant Dis., 92(6):979, June 2008.

Selahattin Danisman, Aalt D J van Dijk, Andrea Bimbo, Froukje van der Wal, Lars Hennig, Stefan de Folter, Gerco C Angenent, and Richard G H Immink. Analysis of functional redundancies within the arabidopsis TCP transcription factor family. J. Exp. Bot., 64(18): 5673–5685, December 2013.

Jean-Michel Daviere and Patrick Achard. A pivotal role of DELLAs in regulating multiple hormone signals. Mol. Plant, 9(1):10–20, January 2016.

Stefan de Folter and Richard G H Immink. Yeast protein-protein interaction assays and screens. Methods Mol. Biol., 754:145–165, 2011.

Amalia Diaz-Granados, Mark G Sterken, Hein Overmars, Roel Ariaans, Martijn Holterman, Somnath S Pokhare, Yulin Yuan, Rikus Pomp, Anna Finkers-Tomczak, Jan Roosien, et al. The effector gprbp-1 of globodera pallida targets a nuclear hect e3 ubiquitin ligase to modulate gene expression in the host. Molecular plant pathology, 21(1):66–82, 2020.

Philip Donkersley, Justine M Blanford, Renan Batista Queiroz, Farley W S Silva, Claudine M Carvalho, Abdullah Mohammed Al-Sadi, and Simon L Elliot. Witch’s broom disease of lime (candidatus phytoplasma aurantifolia): Identifying High-Risk areas by climatic mapping, 2018.

Janani Durairaj, Mehmet Akdel, Dick de Ridder, and Aalt DJ van Dijk. Geometricus represents protein structures as shape-mers derived from moment invariants. Bioinformatics, 36(Supplement 2):i718–i725, 2020.

Robert C Edgar. MUSCLE: a multiple sequence alignment method with reduced time and space complexity. BMC Bioinformatics, 5:113, August 2004.

Nels C Elde and Harmit S Malik. The evolutionary conundrum of pathogen mimicry. Nat. Rev. Microbiol., 7(11):787–797, November 2009.

Sandra K Floyd and John L Bowman. The ancestral developmental tool kit of land plants, 2007.

Jana Fránová, Igor Koloniuk, Ondrej Lenz, and Dimitriyka Sakalieva. Molecular diversity of “candidatus phytoplasma mali” strains associated with apple proliferation disease in bulgarian germplasm collection. Folia Microbiol., 64(3):373–382, May 2019.

K E Frost, P D Esker, R Van Haren, L Kotolski, and R L Groves. Seasonal patterns of aster leafhopper (hemiptera: Cicadellidae) abundance and aster yellows phytoplasma infectivity in wisconsin carrot fields. Environ. Entomol., 42(3):491–502, June 2013.

Christian Gehl, Rainer Waadt, Jorg Kudla, Ralf-R Mendel, and Robert Hansch. New gateway vectors for high throughput analyses of protein–protein interactions by bimolecular fluorescence complementation. Molecular plant, 2(5):1051–1058, 2009.

Eduardo Gonzalez-Grandıo and Pilar Cubas. TCP transcription factors: Evolution, structure, and biochemical function, 2016.

David E Gordon, Joseph Hiatt, Mehdi Bouhaddou, Veronica V Rezelj, Svenja Ulferts, Hannes Braberg, Alexander S Jureka, Kirsten Obernier, Jeffrey Z Guo, Jyoti Batra, et al. Comparative host-coronavirus protein interaction networks reveal pan-viral disease mechanisms. Science, 370(6521):eabe9403, 2020.

Stéphane Guindon, Jean-François Dufayard, Vincent Lefort, Maria Anisimova, Wim Hordijk, and Olivier Gascuel. New algorithms and methods to estimate maximumlikelihood phylogenies: assessing the performance of PhyML 3.0. Syst. Biol., 59(3):307– 321, May 2010.

Geoff M Gurr, Anne C Johnson, Gavin J Ash, Bree A L Wilson, Mark M Ero, Carmel A Pilotti, Charles F Dewhurst, and Minsheng S You. Coconut lethal yellowing diseases: A phytoplasma threat to palms of global economic and social significance, 2016.

S Henikoff and J G Henikoff. Amino acid substitution matrices from protein blocks. Proc. Natl. Acad. Sci. U. S. A., 89(22):10915–10919, November 1992.

Padmini Herath, Gregory A Hoover, Elisa Angelini, and Gary W Moorman. Detection of elm yellows phytoplasma in elms and insects using Real-Time PCR. Plant Dis., 94(11): 1355–1360, November 2010.

Kay Hoffman and Michael D Baron. Boxshade 3.21. pretty printing and shading of multiplealignment files, 1996.

Ayaka Hoshi, Kenro Oshima, Shigeyuki Kakizawa, Yoshiko Ishii, Johji Ozeki, Masayoshi Hashimoto, Ken Komatsu, Satoshi Kagiwada, Yasuyuki Yamaji, and Shigetou Namba. A unique virulence factor for proliferation and dwarfism in plants identified from a phytopathogenic bacterium. Proc. Natl. Acad. Sci. U. S. A., 106(15):6416–6421, April 2009.

Weijie Huang and Saskia Adriane Hogenhout. Interfering with plant developmental timing promotes susceptibility to insect vectors of a bacterial parasite. bioRxiv, pages 2022–03, 2022.

Weijie Huang, Paola Reyes-Caldas, Marina Mann, Shirin Seifbarghi, Alexandra Kahn, Rodrigo PP Almeida, Laure Béven, Michelle Heck, Saskia A Hogenhout, and Gitta Coaker. Bacterial vector-borne plant diseases: unanswered questions and future directions. Molecular plant, 13(10):1379–1393, 2020.

Weijie Huang, Allyson M MacLean, Akiko Sugio, Abbas Maqbool, Marco Busscher, Shu-Ting Cho, Sophien Kamoun, Chih-Horng Kuo, Richard GH Immink, and Saskia A Hogenhout. Parasitic modulation of host development by ubiquitin-independent protein degradation. Cell, 184(20):5201–5214, 2021.

Jaime Huerta-Cepas, François Serra, and Peer Bork. ETE 3: Reconstruction, analysis, and visualization of phylogenomic data. Mol. Biol. Evol., 33(6):1635–1638, June 2016.

Paul Jaccard. THE DISTRIBUTION OF THE FLORA IN THE ALPINE ZONE.1, 1912.

Jinpu Jin, Feng Tian, De-Chang Yang, Yu-Qi Meng, Lei Kong, Jingchu Luo, and Ge Gao. PlantTFDB 4.0: toward a central hub for transcription factors and regulatory interactions in plants, 2017.

J Jović, T Cvrković, M Mitrović, S Krnjajić, A Petrović, M G Redinbaugh, R C Pratt, S A Hogenhout, and I Tosevski. Stolbur phytoplasma transmission to maize by reptalus panzeri and the disease cycle of maize redness in serbia. Phytopathology, 99(9):1053–1061, September 2009.

John Jumper, Richard Evans, Alexander Pritzel, Tim Green, Michael Figurnov, Olaf Ronneberger, Kathryn Tunyasuvunakool, Russ Bates, Augustin Zıdek, Anna Potapenko, et al. Highly accurate protein structure prediction with alphafold. Nature, 596(7873):583–589, 2021.

Mansour Karimi, Dirk Inzé, and Ann Depicker. GATEWAY™ vectors for agrobacteriummediated plant transformation, 2002.

D V Klopfenstein, Liangsheng Zhang, Brent S Pedersen, Fidel Ramírez, Alex War-wick Vesztrocy, Aurélien Naldi, Christopher J Mungall, Jeffrey M Yunes, Olga Botvinnik, Mark Weigel, Will Dampier, Christophe Dessimoz, Patrick Flick, and Haibao Tang. GOATOOLS: A python library for gene ontology analyses. Sci. Rep., 8(1):10872, July 2018.

Sudhir Kumar, Masatoshi Nei, Joel Dudley, and Koichiro Tamura. MEGA: a biologist-centric software for evolutionary analysis of DNA and protein sequences. Brief. Bioinform., 9(4): 299–306, July 2008.

Ruth Le Fevre, Edouard Evangelisti, Thomas Rey, and Sebastian Schornack. Modulation of host cell biology by plant pathogenic microbes. Annu. Rev. Cell Dev. Biol., 31:201–229, September 2015.

Shutian Li. The arabidopsis thaliana TCP transcription factors: A broadening horizon beyond development. Plant Signal. Behav., 10(7):e1044192, 2015.

Qun Liu, Abbas Maqbool, Federico Gabriel Mirkin, Yeshveer Singh, Clare EM Stevenson, David M Lawson, Sophien Kamoun, Weijie Huang, and Saskia Adriane Hogenhout. Bimodular architecture of bacterial effector sap05 drives ubiquitin-independent targeted protein degradation. bioRxiv, pages 2023–06, 2023.

Jessica A Lopez, Yali Sun, Peter B Blair, and M Shahid Mukhtar. TCP three-way handshake: linking developmental processes with plant immunity. Trends Plant Sci., 20(4):238–245, April 2015.

Allyson M MacLean, Akiko Sugio, Olga V Makarova, Kim C Findlay, Victoria M Grieve, Réka Tóth, Mogens Nicolaisen, and Saskia A Hogenhout. Phytoplasma effector SAP54 induces indeterminate Leaf-Like flower development in arabidopsis plants, 2011.

Allyson M MacLean, Zigmunds Orlovskis, Krissana Kowitwanich, Anna M Zdziarska, Gerco C Angenent, Richard G H Immink, and Saskia A Hogenhout. Phytoplasma effector SAP54 hijacks plant reproduction by degrading MADS-box proteins and promotes insect colonization in a RAD23-Dependent manner, 2014.

Sylvie Malembic-Maher, Delphine Desqué, Dima Khalil, Pascal Salar, Bernard Bergey, JeanLuc Danet, Sybille Duret, Marie-Pierre Dubrana-Ourabah, Laure Beven, Ibolya Ember, Zoltan Acs, Michele Della Bartola, Alberto Materazzi, Luisa Filippin, Slobodan Krnjajic, Oliver Krstić, Ivo Toševski, Friederike Lang, Barbara Jarausch, Maria Kölber, Jelena Jović, Elisa Angelini, Nathalie Arricau-Bouvery, Michael Maixner, and Xavier Foissac. When a palearctic bacterium meets a nearctic insect vector: Genetic and ecological insights into the emergence of the grapevine flavescence dorée epidemics in europe. PLoS Pathog., 16 (3):e1007967, March 2020.

Mar Martín-Trillo and Pilar Cubas. TCP genes: a family snapshot ten years later. Trends Plant Sci., 15(1):31–39, January 2010.

Seema Mattoo, Yvonne M Lee, and Jack E Dixon. Interactions of bacterial effector proteins with host proteins, 2007.

Nami Minato, Misako Himeno, Ayaka Hoshi, Kensaku Maejima, Ken Komatsu, Yumiko Takebayashi, Hiroyuki Kasahara, Akira Yusa, Yasuyuki Yamaji, Kenro Oshima, Yuji Kamiya, and Shigetou Namba. The phytoplasmal virulence factor TENGU causes plant sterility by downregulating of the jasmonic acid and auxin pathways. Sci. Rep., 4:7399, December 2014.

M S Mukhtar, A-R Carvunis, M Dreze, P Epple, J Steinbrenner, J Moore, M Tasan, M Galli, T Hao, M T Nishimura, S J Pevzner, S E Donovan, L Ghamsari, B Santhanam, V Romero, M M Poulin, F Gebreab, B J Gutierrez, S Tam, D Monachello, M Boxem, C J Harbort, N McDonald, L Gai, H Chen, Y He, J Vandenhaute, F P Roth, D E Hill, J R Ecker, M Vidal, J Beynon, P Braun, J L Dangl, and European Union Effectoromics Consortium. Independently evolved virulence effectors converge onto hubs in a plant immune system network, 2011.

Olivier Navaud, Patrick Dabos, Elodie Carnus, Dominique Tremousaygue, and Christine Hervé. TCP transcription factors predate the emergence of land plants. J. Mol. Evol., 65 (1):23–33, July 2007.

Sandra Orchard, Mais Ammari, Bruno Aranda, Lionel Breuza, Leonardo Briganti, Fiona Broackes-Carter, Nancy H Campbell, Gayatri Chavali, Carol Chen, Noemi del Toro, Margaret Duesbury, Marine Dumousseau, Eugenia Galeota, Ursula Hinz, Marta Iannuccelli, Sruthi Jagannathan, Rafael Jimenez, Jyoti Khadake, Astrid Lagreid, Luana Licata, Ruth C Lovering, Birgit Meldal, Anna N Melidoni, Mila Milagros, Daniele Peluso, Livia Perfetto, Pablo Porras, Arathi Raghunath, Sylvie Ricard-Blum, Bernd Roechert, Andre Stutz, Michael Tognolli, Kim van Roey, Gianni Cesareni, and Henning Hermjakob. The MIntAct project—IntAct as a common curation platform for 11 molecular interaction databases, 2014.

Zigmunds Orlovskis and Saskia A Hogenhout. A bacterial parasite effector mediates insect vector attraction in host plants independently of developmental changes, 2016.

Zigmunds Orlovskis, Maria Cristina Canale, Mindia Haryono, João Roberto Spotti Lopes, Chih-Horng Kuo, and Saskia A Hogenhout. A few sequence polymorphisms among isolates of maize bushy stunt phytoplasma associate with organ proliferation symptoms of infected maize plants. Ann. Bot., 119(5):869–884, March 2017.

Kenro Oshima, Yoshiko Ishii, Shigeyuki Kakizawa, Kyoko Sugawara, Yutaro Neriya, Misako Himeno, Nami Minato, Chihiro Miura, Takuya Shiraishi, Yasuyuki Yamaji, and Shigetou Namba. Dramatic transcriptional changes in an intracellular parasite enable host switching between plant and insect, 2011.

Jeongmoo Park, Khoa Thi Nguyen, Eunae Park, Jong-Seong Jeon, and Giltsu Choi. Della proteins and their interacting ring finger proteins repress gibberellin responses by binding to the promoters of a subset of gibberellin-responsive genes in arabidopsis. The Plant Cell, 25(3):927–943, 2013.

Pascal Pecher, Gabriele Moro, Maria Cristina Canale, Sylvain Capdevielle, Archana Singh, Allyson MacLean, Akiko Sugio, Chih-Horng Kuo, Joao R S Lopes, and Saskia A Hogenhout. Phytoplasma SAP11 effector destabilization of TCP transcription factors differentially impact development and defence of arabidopsis versus maize. PLoS Pathog., 15(9): e1008035, September 2019.

Paulino Pérez-Rodŕıguez, Diego Mauricio Riaño-Pachón, Luiz Gustavo Guedes Corrêa, Stefan A Rensing, Birgit Kersten, and Bernd Mueller-Roeber. PlnTFDB: updated content and new features of the plant transcription factor database. Nucleic Acids Res., 38(Database issue):D822–7, January 2010.

Efsa Panel on Plant Health (PLH), EFSA Panel on Plant Health (PLH), Claude Bragard, Katharina Dehnen-Schmutz, Paolo Gonthier, Josep Anton Jaques Miret, Annemarie Fejer Justesen, Alan MacLeod, Christer Sven Magnusson, Panagiotis Milonas, Juan A Navas-Cortes, Stephen Parnell, Roel Potting, Philippe Lucien Reignault, Hanshermann Thulke, Wopke Van der Werf, Antonio Vicent Civera, Jonathan Yuen, Lucia Zappalà, Domenico Bosco, Michela Chiumenti, Francesco Di Serio, Luciana Galetto, Cristina Marzachí, Marco Pautasso, and Marie-agnès Jacques. Pest categorisation of the non-EU phytoplasmas of cydonia mill., fragaria l., malus mill., prunus l., pyrus l., ribes l., rubus l. and vitis L, 2020.

Jose L Pruneda-Paz, Ghislain Breton, Dawn H Nagel, S Earl Kang, Katia Bonaldi, Colleen J Doherty, Stephanie Ravelo, Mary Galli, Joseph R Ecker, and Steve A Kay. A genome-scale resource for the functional characterization of arabidopsis transcription factors. Cell Rep., 8(2):622–632, July 2014.

A Rambaut. FigTree v1. 4.0. a graphical viewer of phylogenetic trees, 2012.

Pratibha Ravindran and Prakash P Kumar. Regulation of seed germination: The involvement of multiple forces exerted via gibberellic acid signaling. Mol. Plant, 12(10):1416–1417, October 2019.

Patricia A Rodriguez, Michael Rothballer, Soumitra Paul Chowdhury, Thomas Nussbaumer, Caroline Gutjahr, and Pascal Falter-Braun. Systems biology of Plant-Microbiome interactions, 2019.

B A Roy. Floral mimicry by a plant pathogen, 1993.

Florian Rümpler, Lydia Gramzow, Günter Theißen, and Rainer Melzer. Did convergent protein evolution enable phytoplasmas to generate ‘zombie plants’?, 2015.

M E Santos-Cervantes, J A Chávez-Medina, J Méndez-Lozano, and N E Leyva-López. Detection and molecular characterization of two little leaf phytoplasma strains associated with pepper and tomato diseases in guanajuato and sinaloa, mexico. Plant Dis., 92(7): 1007–1011, July 2008.

Skipper Seabold and Josef Perktold. statsmodels: Econometric and statistical modeling with python. In 9th Python in Science Conference, 2010.

Edouard Severing, Luigi Faino, Suraj Jamge, Marco Busscher, Yang Kuijer-Zhang, Francesca Bellinazzo, Jacqueline Busscher-Lange, Virginia Fernández, Gerco C Angenent, Richard GH Immink, et al. Arabidopsis thaliana ambient temperature responsive lncrnas. BMC plant biology, 18(1):1–10, 2018.

Fabian Sievers, Andreas Wilm, David Dineen, Toby J Gibson, Kevin Karplus, Weizhong Li, Rodrigo Lopez, Hamish McWilliam, Michael Remmert, Johannes Söding, Julie D Thompson, and Desmond G Higgins. Fast, scalable generation of high-quality protein multiple sequence alignments using clustal omega. Mol. Syst. Biol., 7:539, October 2011.

W A Sinclair, M L Gleason, H M Griffiths, J K Iles, N Zriba, D V Charlson, J C Batzer, and T H Whitlow. Responses of 11 fraxinus cultivars to ash yellows phytoplasma strains of differing aggressiveness. Plant Dis., 84(7), July 2000.

Greg Slodkowicz and Nick Goldman. Integrated structural and evolutionary analysis reveals common mechanisms underlying adaptive evolution in mammals, 2020.

Evelyn Strauss. Microbiology. phytoplasma research begins to bloom. Science, 325(5939): 388–390, July 2009.

Akiko Sugio, Heather N Kingdom, Allyson M MacLean, Victoria M Grieve, and Saskia A Hogenhout. Phytoplasma protein effector SAP11 enhances insect vector reproduction by manipulating plant development and defense hormone biosynthesis. Proc. Natl. Acad. Sci. U. S. A., 108(48):E1254–63, November 2011a.

Akiko Sugio, Allyson M MacLean, Heather N Kingdom, Victoria M Grieve, R Manimekalai, and Saskia A Hogenhout. Diverse targets of phytoplasma effectors: From plant development to defense against insects, 2011b.

Akiko Sugio, Allyson M MacLean, and Saskia A Hogenhout. The small phytoplasma virulence effector SAP11 contains distinct domains required for nuclear targeting and CIN-TCP binding and destabilization, 2014.

Xudong Sun, Zhenhua Feng, Laisheng Meng, Jian Zhu, and Anja Geitmann. Arabidopsis ASL11/LBD15 is involved in shoot apical meristem development and regulates WUS expression. Planta, 237(5):1367–1378, May 2013.

Lokesh P Tripathi, Yi-An Chen, Kenji Mizuguchi, and Eiji Morita. Network-Based analysis of Host-Pathogen interactions, 2019.

Leigh Van Valen. A new evolutionary law. Evolutionary Theory, 1(1):1–30, 1973.

Allegra Via, Bora Uyar, Christine Brun, and Andreas Zanzoni. How pathogens use linear motifs to perturb host cell networks. Trends Biochem. Sci., 40(1):36–48, January 2015.

Pauli Virtanen, Ralf Gommers, Travis E Oliphant, Matt Haberland, Tyler Reddy, David Cournapeau, Evgeni Burovski, Pearu Peterson, Warren Weckesser, Jonathan Bright, Stéfan J van der Walt, Matthew Brett, Joshua Wilson, K Jarrod Millman, Nikolay Mayorov, Andrew R J Nelson, Eric Jones, Robert Kern, Eric Larson, C J Carey, Ilhan Polat, Yu Feng, Eric W Moore, Jake VanderPlas, Denis Laxalde, Josef Perktold, Robert Cimrman, Ian Henriksen, E A Quintero, Charles R Harris, Anne M Archibald, Antônio H Ribeiro, Fabian Pedregosa, Paul van Mulbregt, and SciPy 1.0 Contributors. SciPy 1.0: fundamental algorithms for scientific computing in python. Nat. Methods, 17(3):261–272, March 2020.

Nan Wang, Haizhen Yang, Zhiyuan Yin, Wenting Liu, Liying Sun, and Yunfeng Wu. Phytoplasma effector SWP1 induces witches’ broom symptom by destabilizing the TCP transcription factor BRANCHED1, 2018.

Wei Wei, Robert E Davis, Gary R Bauchan, and Yan Zhao. New symptoms identified in phytoplasma-infected plants reveal extra stages of pathogen-induced meristem fatederailment. Molecular Plant-Microbe Interactions, 32(10):1314–1323, 2019.

Phyllis G Weintraub and Leann Beanland. Insect vectors of phytoplasmas. Annu. Rev. Entomol., 51:91–111, 2006.

Ralf Weßling, Petra Epple, Stefan Altmann, Yijian He, Li Yang, Stefan R Henz, Nathan Mc-Donald, Kristin Wiley, Kai Christian Bader, Christine Gläßer, M Shahid Mukhtar, Sabine Haigis, Lila Ghamsari, Amber E Stephens, Joseph R Ecker, Marc Vidal, Jonathan D G Jones, Klaus F X Mayer, Emiel Ver Loren van Themaat, Detlef Weigel, Paul Schulze-Lefert, Jeffery L Dangl, Ralph Panstruga, and Pascal Braun. Convergent targeting of a common host Protein-Network by pathogen effectors from three kingdoms of life, 2014.

Jinrui Xu and Yang Zhang. How significant is a protein structure similarity with tm-score= 0.5? Bioinformatics, 26(7):889–895, 2010.

Li Yang, Paulo José Pereira, Surojit Biswas, Omri M Finkel, Yijian He, Isai Salas-Gonzalez, Marie E English, Petra Epple, Piotr Mieczkowski, and Jeffery L Dangl. Pseudomonas syringae type III effector HopBB1 promotes host transcriptional repressor degradation to regulate phytohormone responses and virulence, 2017.

Yukiko Yasui, Keiko Mukougawa, Mitsuhiro Uemoto, Akira Yokofuji, Ryota Suzuri, Aiko Nishitani, and Takayuki Kohchi. The Phytochrome-Interacting VASCULAR PLANT ONE– ZINC FINGER1 and VOZ2 redundantly regulate flowering in arabidopsis, 2012.

Yang Zhang and Jeffrey Skolnick. Tm-align: a protein structure alignment algorithm based on the tm-score. Nucleic acids research, 33(7):2302–2309, 2005.

