## Supplementary information for "Protein interaction mapping reveals widespread targeting of development-related host transcription factors by phytoplasma effectors"

### 7 Supplementary Information

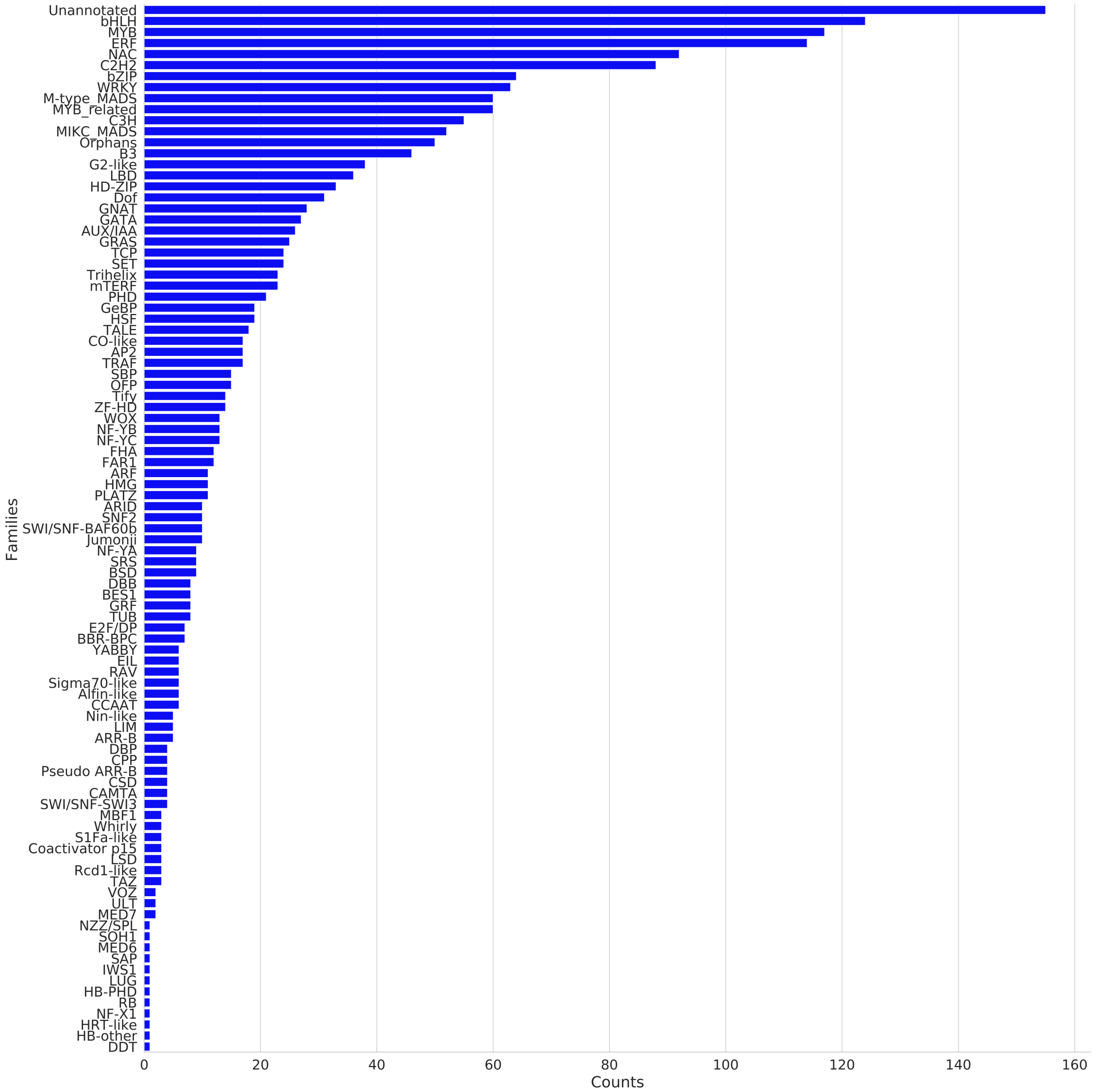


Figure S1: Distribution of host protein families in the screened library.

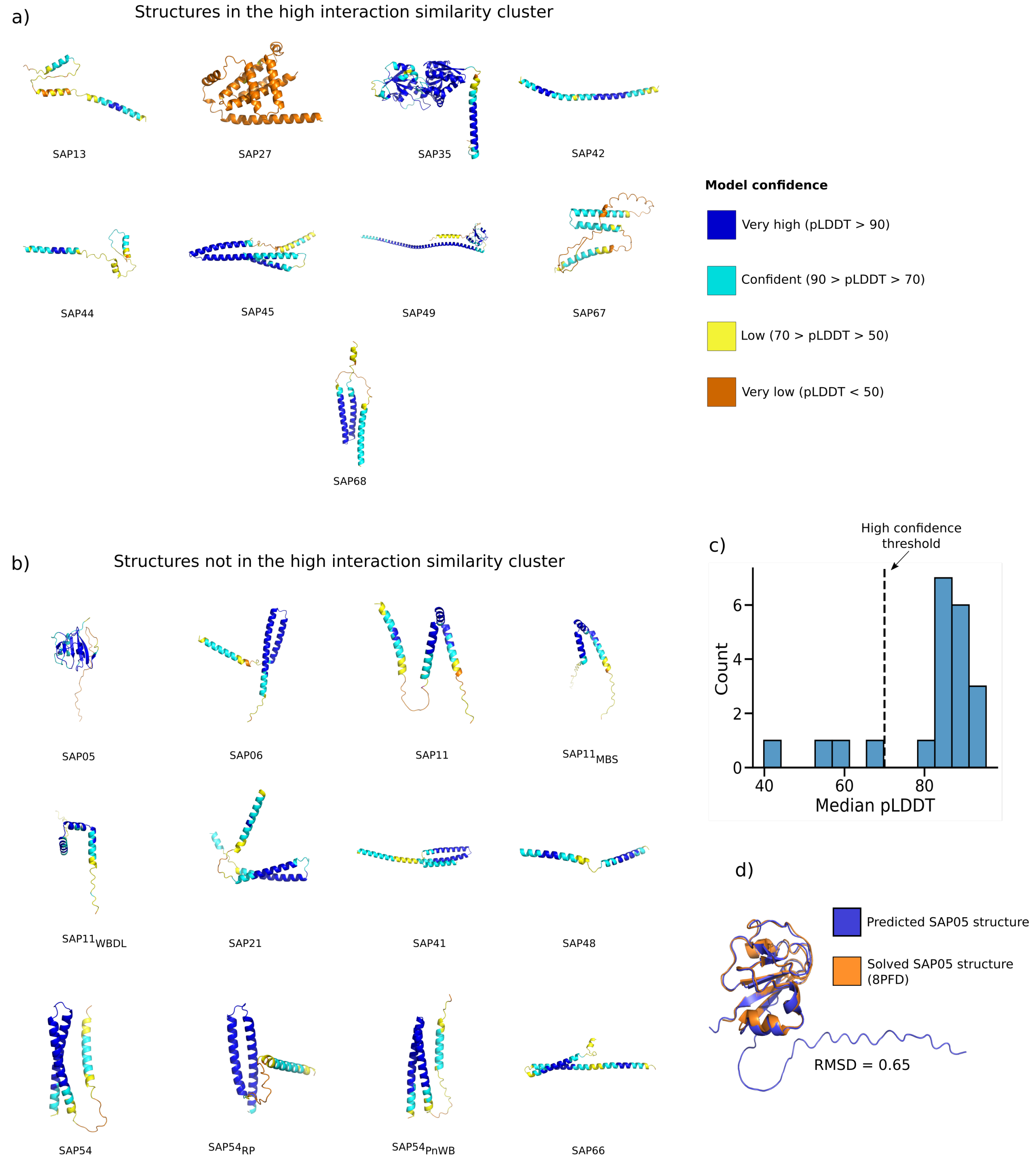


Figure S2: AlphaFold2-predicted structures of phytoplasma effectors in our dataset. Structures are coloured according to their predicted local Distance Difference Test (pLDDT), a per-residue metric of predicted accuracy. (a) Predicted structures for effectors in the high interaction similarity cluster. (b) Predicted structures for effectors not in the cluster. (c) Histogram of overall predicted structure quality, expressed as median pLDDT per protein. (d) Superposition of the predicted structure of SAP05 and its experimentally solved counterpart.

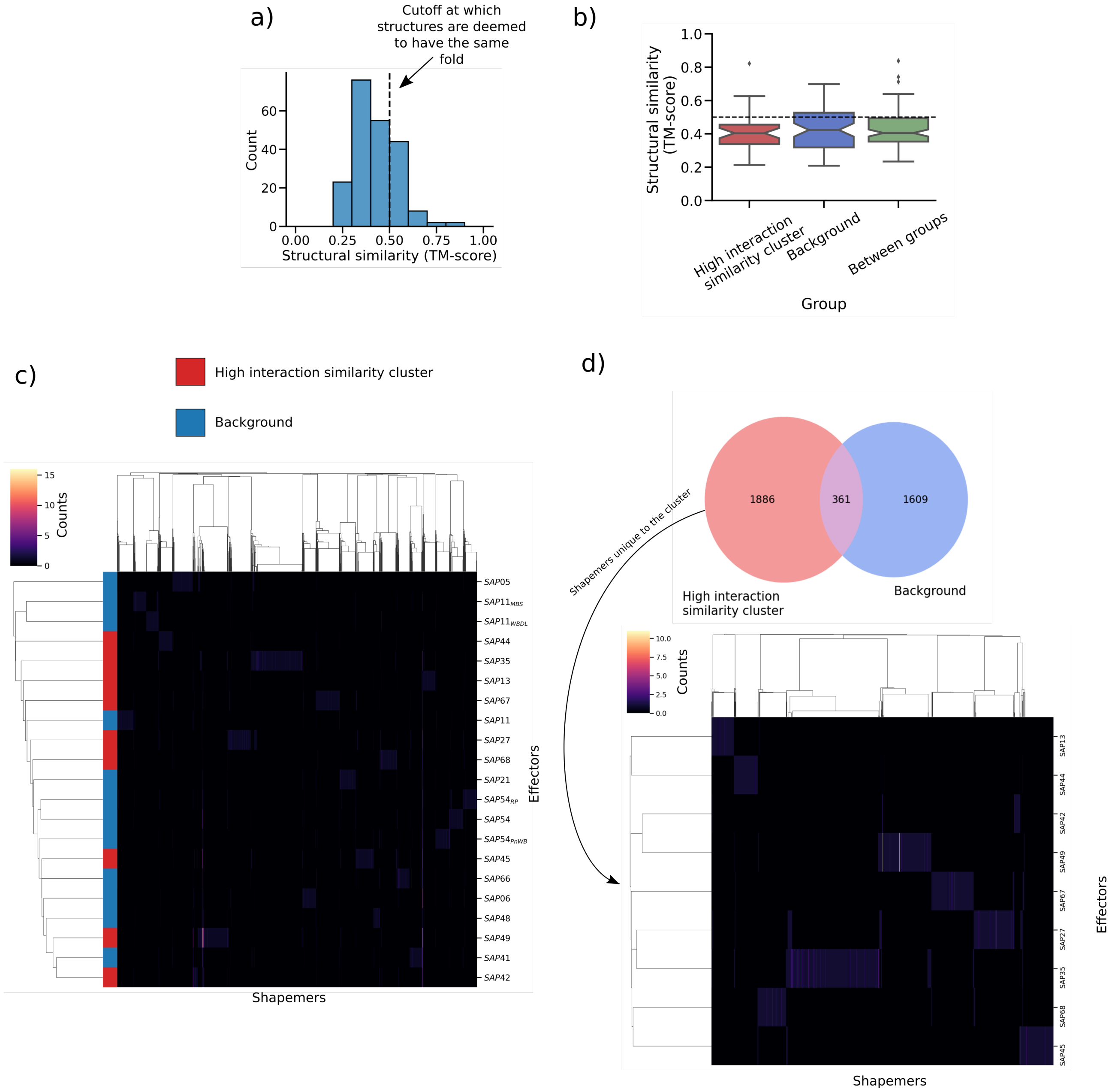


Figure S3: Global and local comparison of the predicted structures of phytoplasma effectors. (a) Histogram of global pairwise structural similarities, quantified as TM-scores. The vertical dashed line indicates the empirically determined threshold of 0.5 at which proteins often share the same fold. (b) Pairwise structural similarities on different subsets of phytoplasma effectors: proteins within the high interaction similarity cluster, proteins not in this cluster (considered as the background), and comparisons between the two groups. Notches on the boxplots indicate 95% confidence intervals computed using 1000 bootstrap resamples. (c) Clustermap of all phytoplasma effectors according to their structural embedding using cosine distance as a similarity metric. Red rows indicate effectors within the high interaction similarity cluster; blue rows indicate effectors in the background. (d) Panel above: Venn diagram of shapemers in the high interaction similarity cluster and the background. Panel below: clustermap of effectors within the high interaction similarity cluster, using only shapemers found exclusively within this set of proteins.

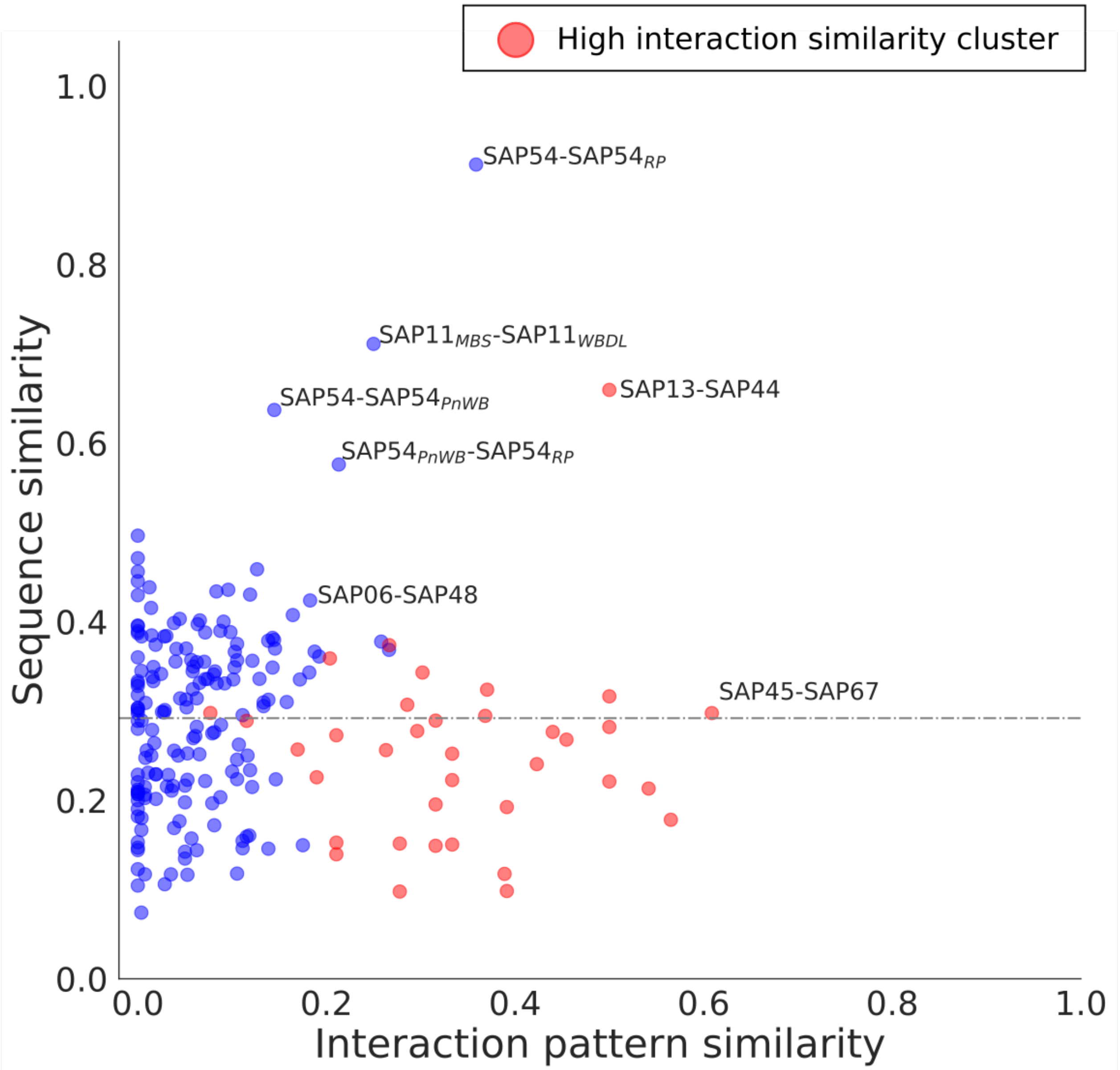


Figure S4: Relationship between effector interaction similarity and sequence similarity. Effectors belonging to the cluster of sequences with similar interaction patterns are marked in red; outliers and interesting examples are annotated with the pair they represent. The horizontal dashed line indicates median sequence similarity.

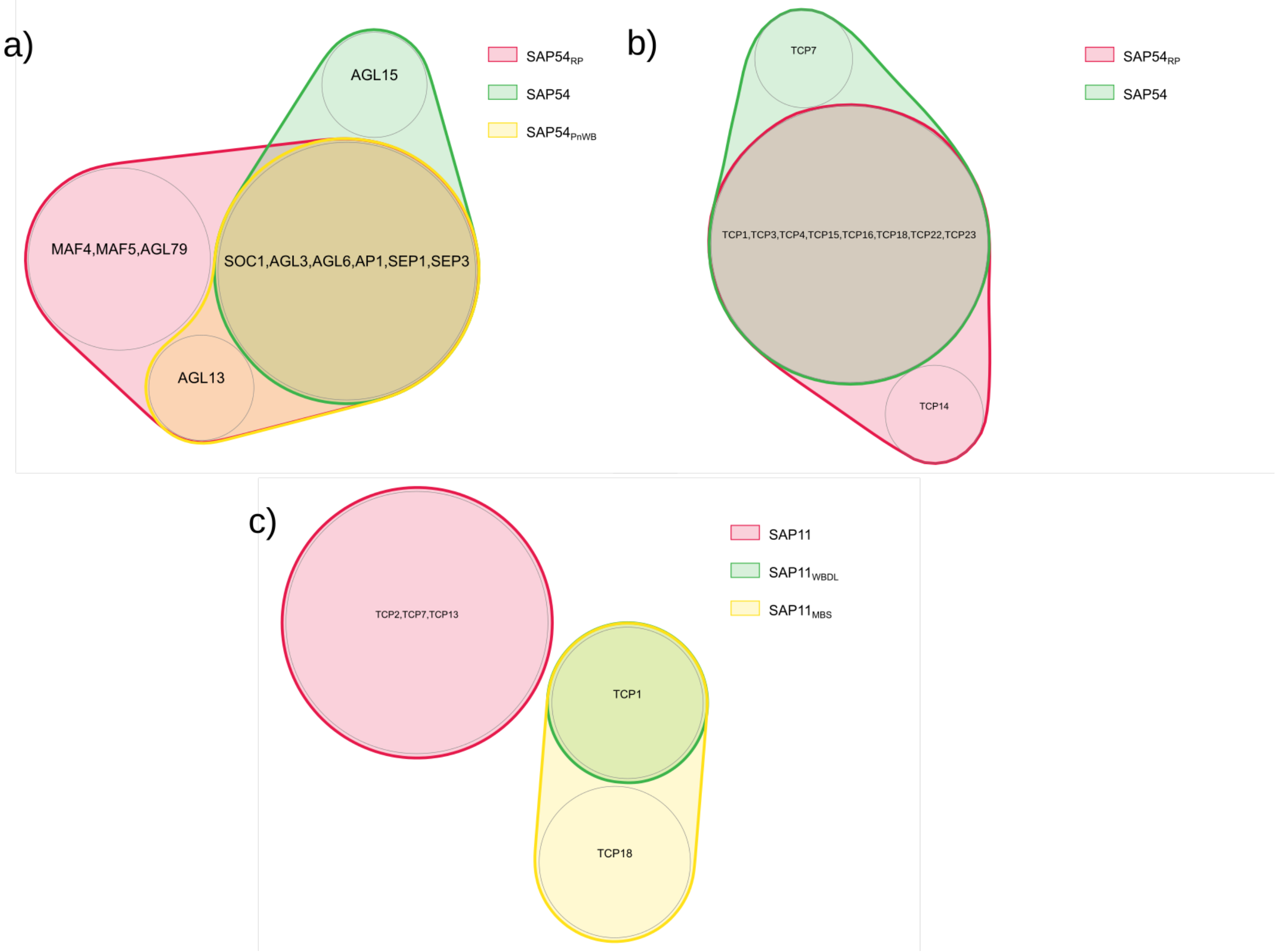


Figure S5: Venn diagrams of interactions across different orthologous SAPs. (a) Interactions between different SAP54 orthologs and MIKC MADS-box proteins. (b) Interactions between different SAP54 orthologs and TCP transcription factors. (c) Interactions between different SAP11 orthologs and TCP transcription factors.

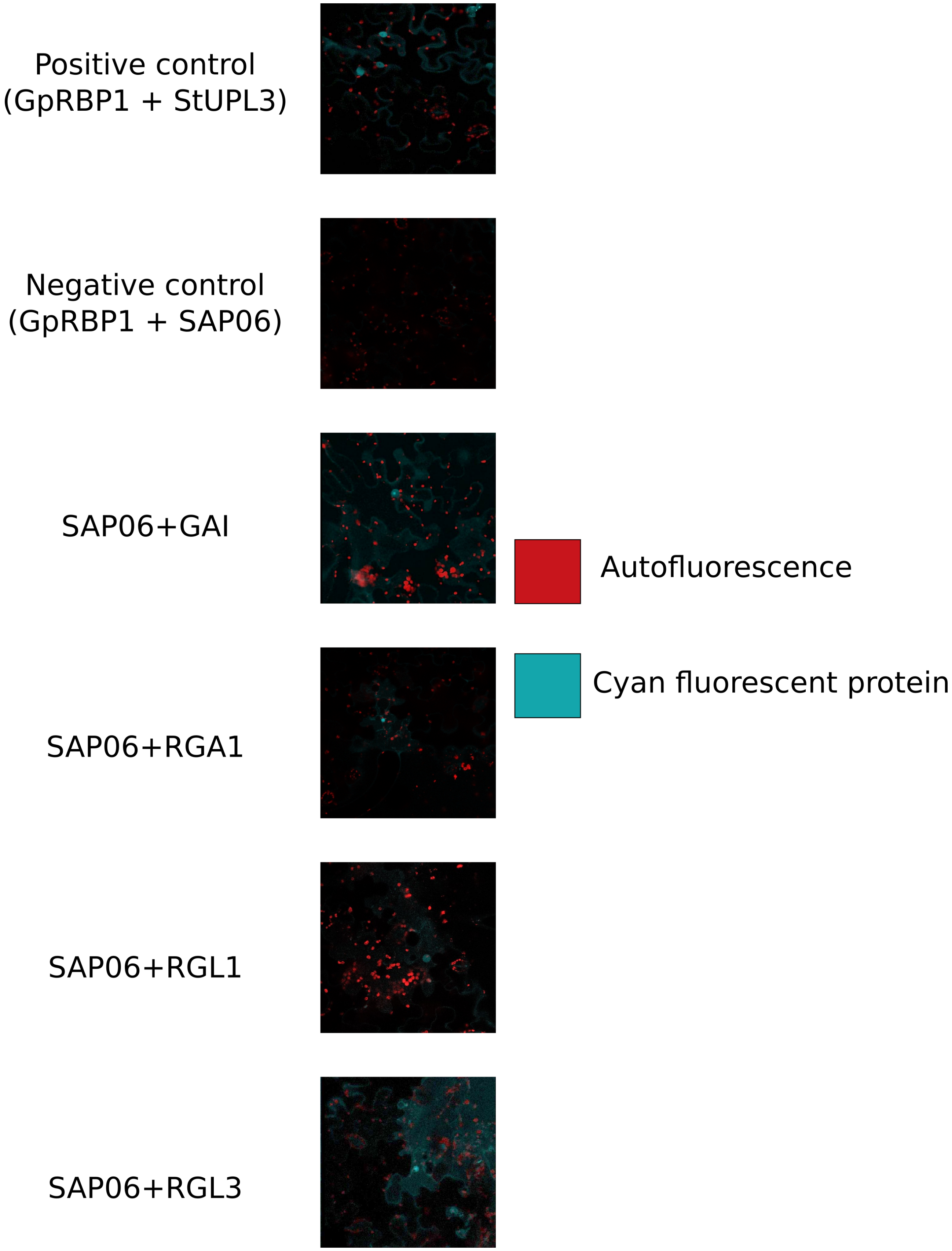


Figure S6: Confirmation by BiFC of interactions between SAP06 and four different DELLA transcription factors. Red signal comes from autofluorescing plastids and chloroplasts; cyan signal is from the reconstituted cyan fluorescent protein (CFP), indicating PPIs. The interaction with SAP06 is confirmed for all four different DELLA proteins tested (GAI, RGA1, RGL1 and RGL3).

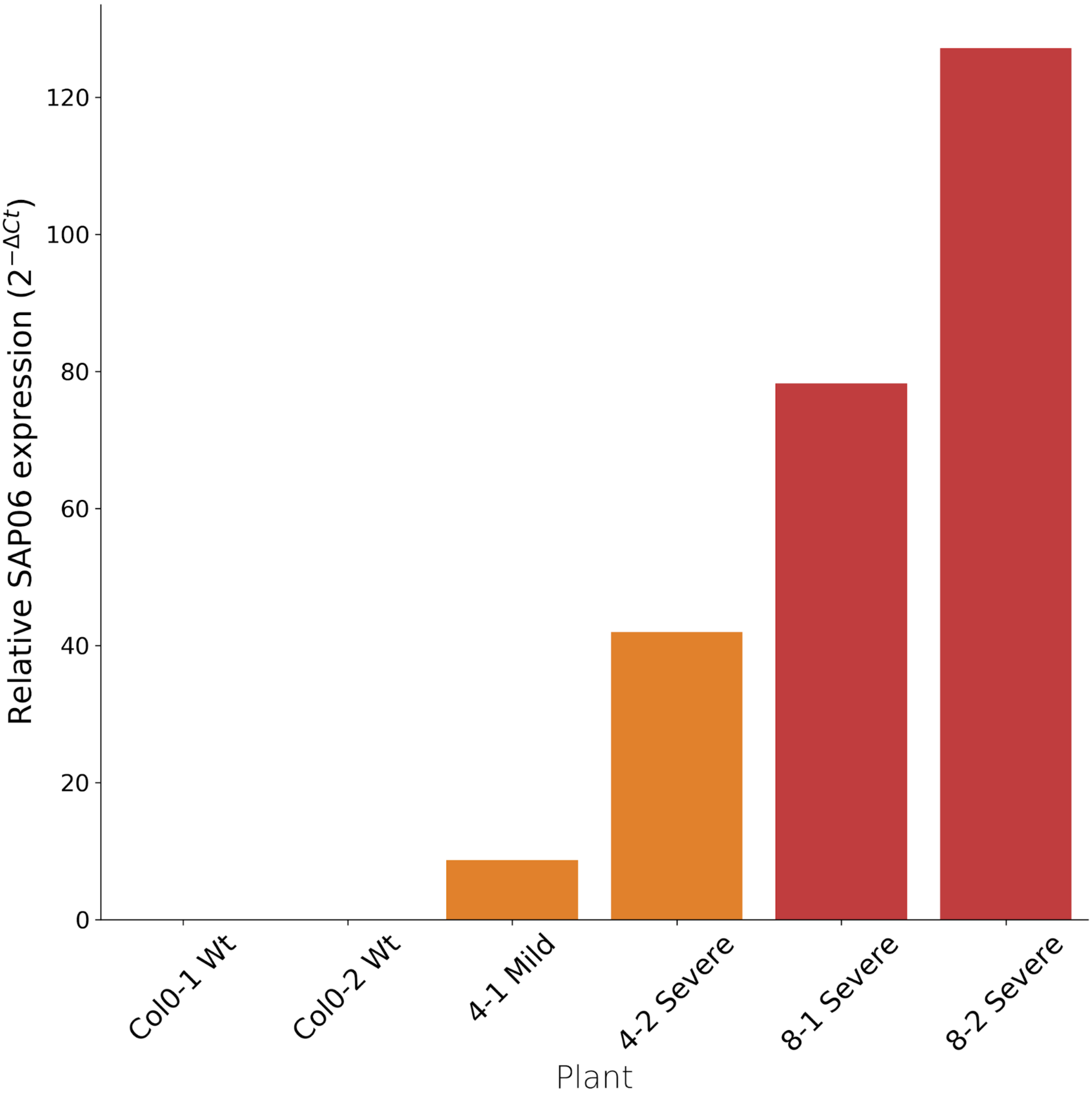


Figure S7: Relative ectopic expression levels of SAP06 in four individual transgenic plants in the progeny of two independent transformed lines, Line #4 (plant 4-1 and plant 4-2) and line #8 (plant 8-1 and plant 8-2). Three out of four individual plants showed a severe overexpression phenotype (strongly reduced in size). Plant 4-1 had a mild phenotype, but clearly deviates from Col-0 wild type plants (Fig. 4a, main text). Two Col-0 plants were included as negative controls.

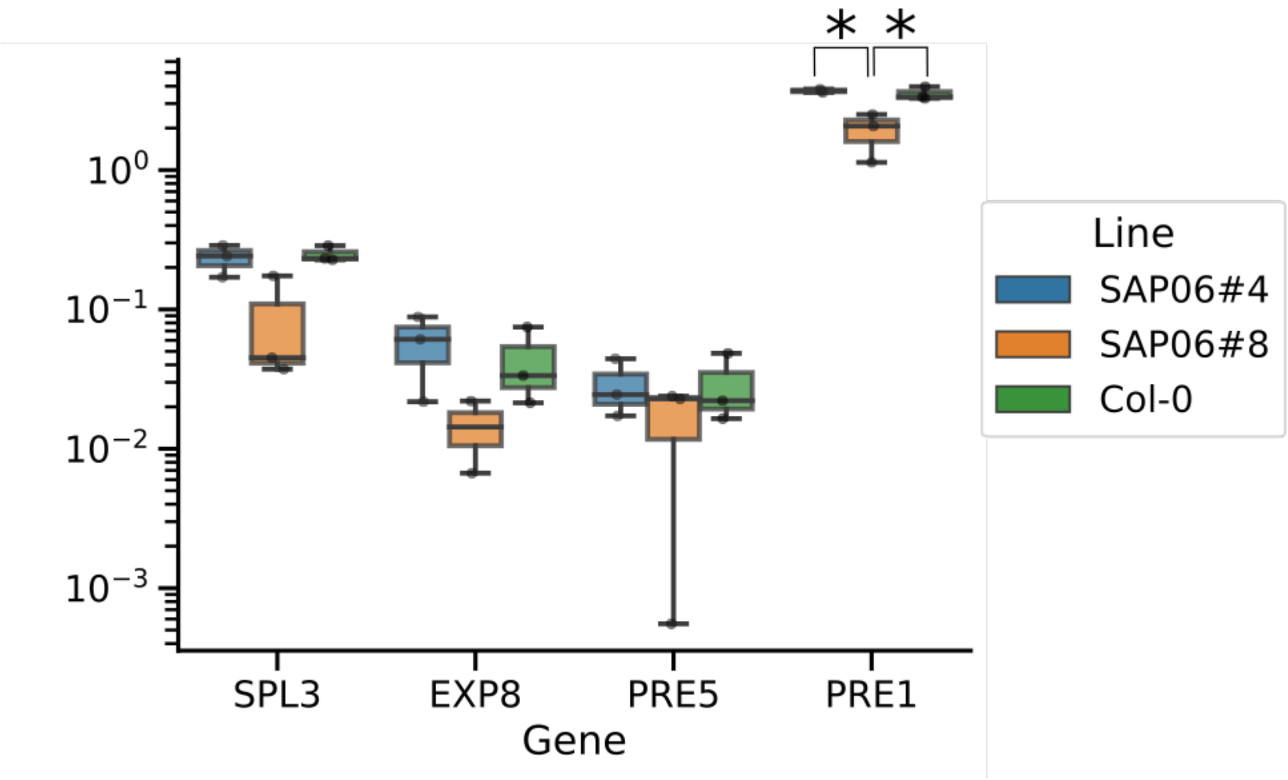


Figure S8: qRT-PCR results measuring expression of four known DELLA target genes in individuals belonging to the SAP06#4, SAP06#8 and Col-0 lines using SAND as a reference gene. Three individuals of each line were used for measurements. * indicates statistical significance (adjusted $p < 0.05$).

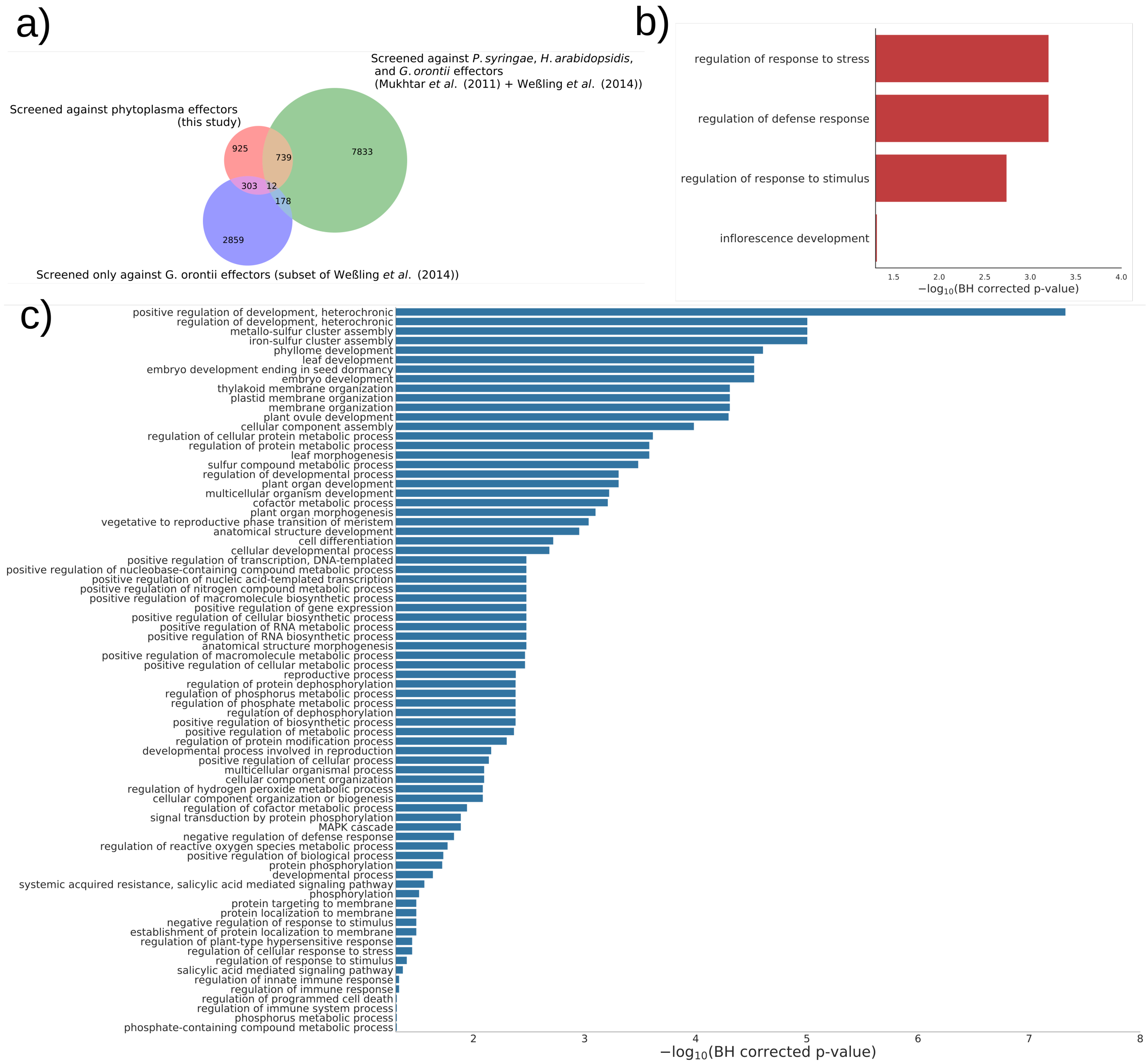


Figure S9: (a) Overlap between host protein libraries used in different yeast-two hybrid assays. (b) Enriched biological process GO terms in host proteins that interact with effectors from all pathogens (including phytoplasma). (c) Enriched biological process GO terms in host proteins that are highly targeted in phytoplasma but do not interact with any effector assayed in *P. syringae*, *H. arabidopsidis* or *G. orontii*. Note that the horizontal axis starts at the statistical significance threshold (0.05).

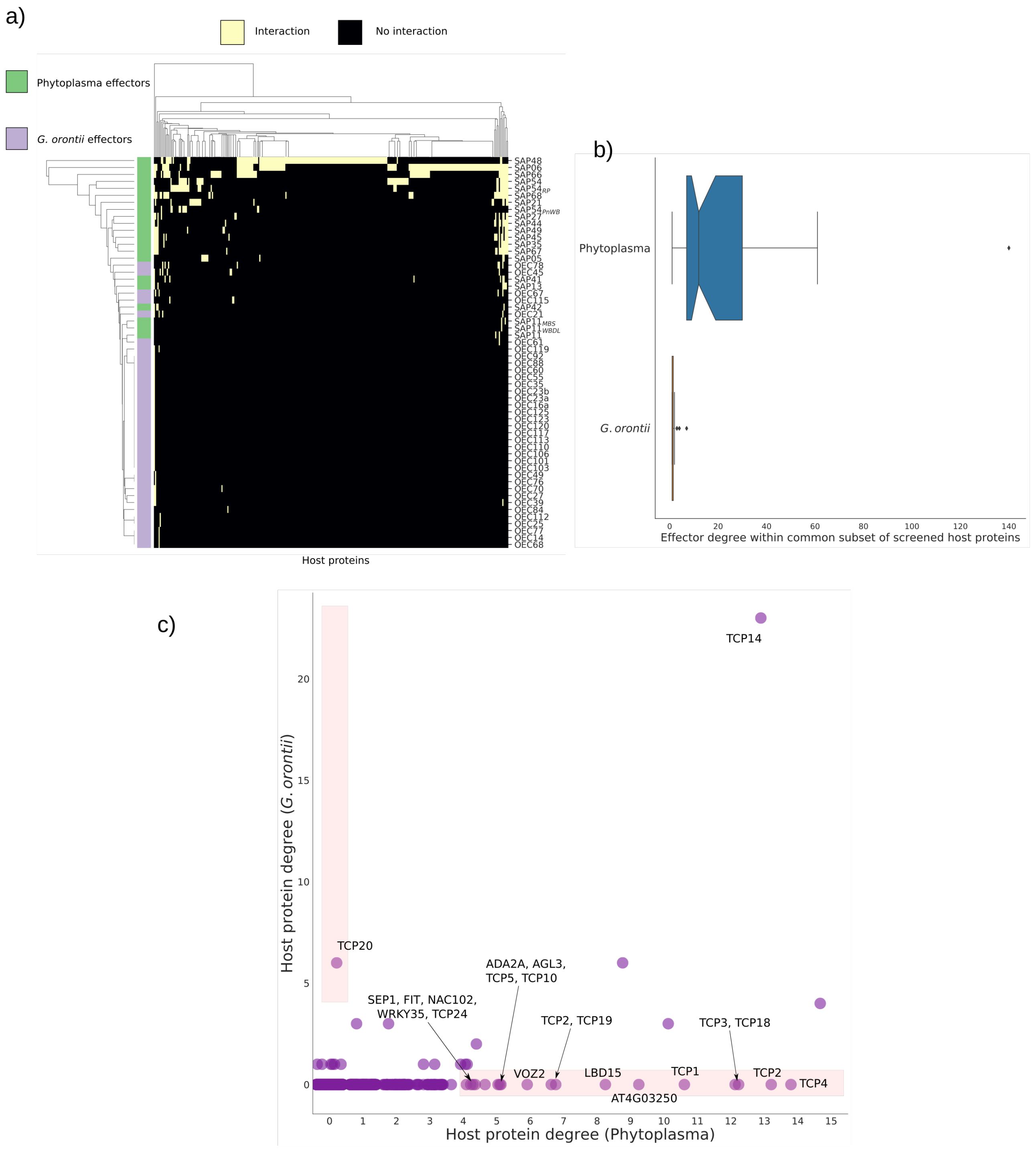


Figure S10: Comparison between phytoplasma-host and *G. orontii*-host PPI networks, using the common subset of host proteins assayed against the effectors. 1056 host proteins were shared between the two sets, 757 of which did not show interactions with any effector. These were removed to calculate interaction pattern similarities. (a) Clustermap of interaction patterns in the two networks. Clustering is done using (1 - Jaccard similarity) as a distance metric. (b) Boxplot of the degrees of effectors. Notches indicate the width of a 95% confidence interval of the median degree, determined by 1000 bootstrap resamples. (c) Comparison of host protein degrees in the two networks. Host proteins that are highly targeted by one pathogen but are not targeted by the other are highlighted.
